# Specific prostaglandins are produced in the migratory cells and the surrounding substrate to promote *Drosophila* border cell migration

**DOI:** 10.1101/2023.06.23.546291

**Authors:** Samuel Q. Mellentine, Anna S. Ramsey, Jie Li, Hunter N. Brown, Tina L. Tootle

**Affiliations:** Anatomy and Cell Biology, University of Iowa Carver College of Medicine, Iowa City, IA 52242

**Keywords:** Border cells, *Drosophila*, cell migration, prostaglandins, PGE_2_, PGF_2α_, integrins, myosin

## Abstract

A key regulator of collective cell migration is prostaglandin (PG) signaling. However, it remains largely unclear whether PGs act within the migratory cells or their microenvironment to promote migration. Here we use *Drosophila* border cell migration as a model to uncover the cell-specific roles of two PGs in collective migration. Prior work shows PG signaling is required for on-time migration and cluster cohesion. We find that the PGE_2_ synthase cPGES is required in the substrate, while the PGF_2α_ synthase Akr1B is required in the border cells for on-time migration. Akr1B acts in both the border cells and their substrate to regulate cluster cohesion. One means by which Akr1B regulates border cell migration is by promoting integrin-based adhesions. Additionally, Akr1B limits myosin activity, and thereby cellular stiffness, in the border cells, whereas cPGES limits myosin activity in both the border cells and their substrate. Together these data reveal that two PGs, PGE_2_ and PGF_2α_, produced in different locations, play key roles in promoting border cell migration. These PGs likely have similar migratory versus microenvironment roles in other collective cell migrations.

## Introduction

Coordinated migration of groups of cells, termed collective cell migration, drives development and tissue repair, and is co-opted during cancer metastasis (Friedl and Gilmour, 2009; Scarpa and Mayor, 2016). Such migrations are regulated by factors from both the migrating cells and their microenvironment (Fife et al., 2014; Kai et al., 2016; Stuelten et al., 2018). One means of regulating collective migration is prostaglandin (PG) signaling (Menter and Dubois, 2012; Tootle, 2013; Kobayashi et al., 2018). PGs are short-range lipid signaling molecules (Funk, 2001; Tootle, 2013). PG signaling begins with cyclooxygenase (COX) enzymes converting arachidonic acid into the prostaglandin precursor (PGH_2_), which is used by PG-type specific synthases to produce bioactive PGs (PGI_2_, PGE_2_, PGF_2α_, PGD_2_ and TXA_2_). These PGs signal in an autocrine or paracrine fashion to activate G-protein coupled receptors (GPCRs) and downstream signaling.

PGs promote collective migration. In zebrafish, global loss of PG signaling impairs migration and delays gastrulation (Cha et al., 2005; Cha et al., 2006). Exogenous application of PGs to cultured cancer cells increases motility. In patients, inhibition of PG synthesis via COX inhibitors reduces the risk of cancer metastasis (Li et al., 2012; Menter and Dubois, 2012). While these studies established PG signaling promotes migration, it remains unclear which PG or PGs regulate migration, whether PGs act within the migrating cells and/or their microenvironment, and how specific PGs drive migration.

To address these questions, we use the *in vivo*, collective migration of the border cells during *Drosophila* oogenesis. Each ovary contains 16-20 ovarioles, chains of sequentially developing follicles (Giedt and Tootle, 2023). There are 14 stages of follicle development and each follicle is comprised of 16 germline cells – 15 nurse cells and one oocyte – and ∼650 somatic cells termed follicle cells. During Stage 9 (S9), 6-8 follicle cells are specified as the border cells, delaminate from the follicular epithelium and migrate posteriorly between the nurse cells to the oocyte (Montell, 2003; Montell et al., 2012). Thus, the nurse cells comprise both the microenvironment and the substrate for border cell migration. Throughout the migration, the border cell cluster is in line with the position of the outer follicle cells, providing an internal control for on-time migration (Fig. 1A).

**Figure 1:**
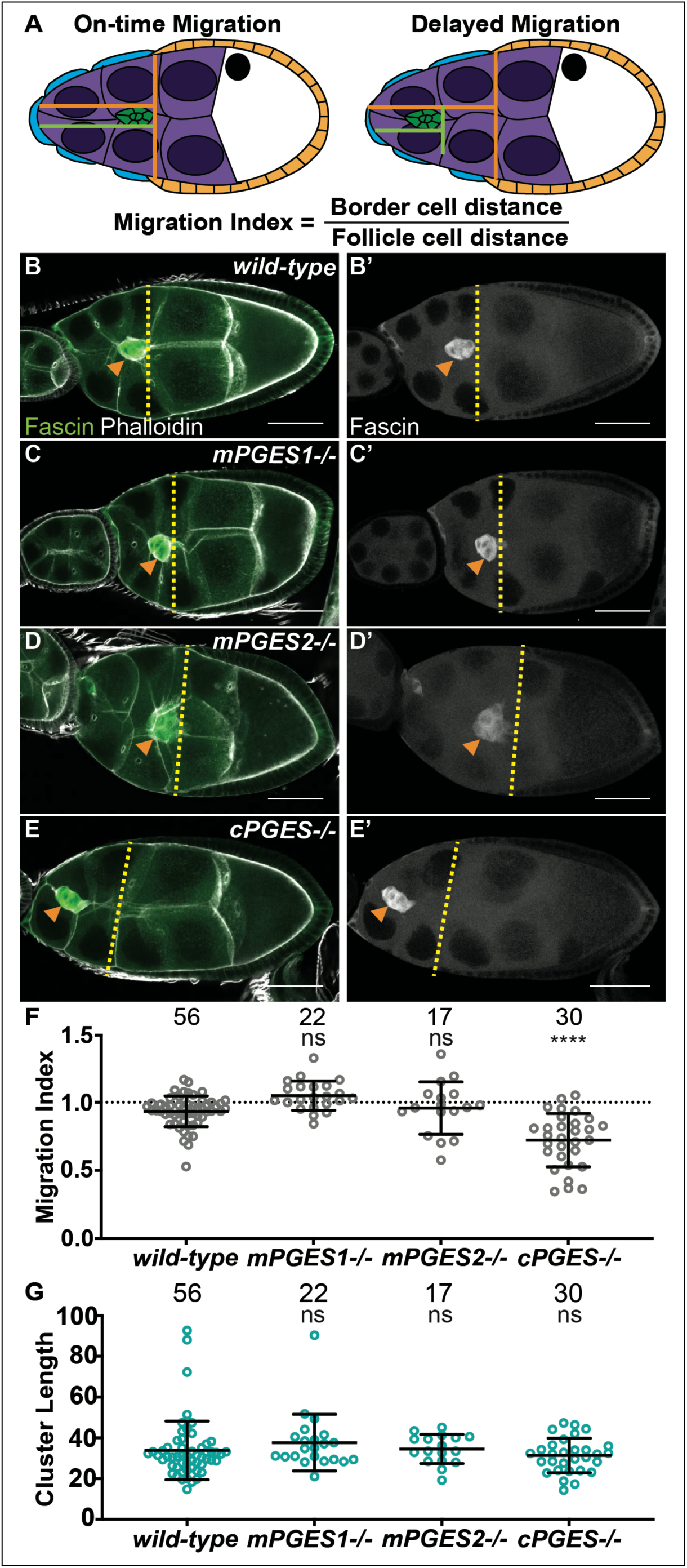
cPGES is required for on-time border cell migration. A. Schematic of on-time (left) and delayed (right) border cell migration and how the migration index is calculated; anterior is to the left and posterior is to the right. The border cell cluster (green), outer follicle cells (orange), stretch follicle cells (blue), nurse cells (purple), and oocyte (white) are diagramed. B-D’. Maximum projections of 3 confocal slices of S9 follicles stained for Fascin (green in merge) and F-actin (phalloidin, white in merge). Orange arrowheads indicate the border cell cluster and yellow dashed lines indicate the position of the outer follicle cells. Images brightened by 30% to increase clarity. Scale bars = 50μm. B-B’. *wild-type* (*yw*). C-C’. *mPGES1*-/- (*mPGES1^KG04713/KG04713^*). D-D’. *mPGES2-/-* (*mPGES2^EY13245/EY13245^*). E-E’. *cPGES-/-* (*cPGES^EY05607/EY05607^*). F-G. Graphs of migration index (F) and border cell cluster length (G) for the indicated genotypes. Circle = single follicle; n = number of follicles. In F, the dotted line indicates on-time border cell migration. For F-G, lines = averages and error bars = SD. ns>0.05, **** p<0.0001, unpaired t-test, two-tailed. In wild-type follicles, throughout S9, the migrating border cell cluster is in-line with the outer follicle cells (A, left). When migration is delayed, the cluster remains anterior to the follicle cells (A, right). We take advantage of this coordination to calculate the migration index (A), which is the distance of the border cell cluster divided by the distance of the outer follicle cells. On-time migration results in a migration index of ∼1, while delayed migration is <1. Like wild-type (B-B’), loss of mPGES1 or mPGES2 exhibit on-time border cell migration (C-D’, F), whereas loss of cPGES delays migration (E-F). Border cell cluster length is unaffected by loss of mPGES1, mPGES2 or cPGES (G).

Border cell migration requires PG signaling (Fox et al., 2020). *Drosophila* have a single COX-like enzyme, Pxt (Tootle and Spradling, 2008); subsequently referred to as dCOX1. Loss of dCOX1 both delays migration and elongates the cluster, indicative of defective cohesion. Cell-specific RNAi experiments reveal strong knockdown of dCOX1 in the border cells delays migration and causes cluster compaction, whereas mild knockdown in the nurse cells (i.e. the substrate) decreases cluster cohesion (Fox et al., 2020). These results indicate PGs are produced in both the migratory cells and their substrate to regulate migration, and PGs from the different cell types may control distinct aspects of border cell migration, on-time migration and cluster cohesion.

We sought to determine which PG or PGs are produced in the border cells and/or the substrate, and the downstream mechanisms whereby they promote on-time migration and maintain cluster cohesion. We find that PGE_2_ produced in the nurse cells by the cytosolic PGE_2_ synthase (cPGES), *Drosophila* p23, is required for on-time migration. Whereas, PGF_2α_ produced in the border cells by Akr1B promotes migration. PGE_2_ signaling does not impact cluster cohesion, but PGF_2α_ signaling has distinct cell-specific roles. Knockdown of Akr1B in the border cells results in compacted clusters, whereas knockdown in the substrate causes cluster elongation. One potential means by which these PGs may promote border cell migration is by regulating integrins. Previously, we found dCOX1 is required for integrin localization to the surface of the border cell cluster (Fox et al., 2020). Here we find that Akr1B is required for integrin localization. Another means of controlling migration is by regulating the balance of forces between the migrating border cells and their substrate, the nurse cells (Majumder et al., 2012; Aranjuez et al., 2016). This mechanoreciprocity is mediated at the cellular level by the activation of non-muscle myosin II (subsequently referred to as myosin). We find that cPGES is required to limit myosin activity in both the border cells and the substrate, whereas Akr1B only limits it within the border cells. These data lead to the model that Akr1B produces PGF_2α_, primarily within the border cells, to regulate both integrin localization and myosin activity within the border cells. In the substrate, cPGES produces PGE_2_, which regulates myosin activity within the substrate and the border cells. Ultimately, the synthesis of both PGE_2_ and PGF_2α_ is required for on-time border cell migration, while only PGF_2α_ modulates cluster cohesion. This work provides the first evidence that multiple types of PGs, produced from different cellular sources, work in concert to control collective cell migration by both overlapping and distinct mechanisms. Given the conservation of PG signaling, such a multi-cellular and multi-PG mechanism of promoting cell migration is likely conserved across organisms and tissues.

## Results

### cPGES is required for on-time border cell migration

Loss of all PG synthesis delays border cell migration and elongates clusters (Fox et al., 2020), however, which specific PGs are involved remains unknown. We first assessed the role of PGE_2_. There are three PGE_2_ synthases, microsomal PGES1 (mPGES1), mPGES2, and cPGES (Jakobsson et al., 1999; Tanioka et al., 2000; Tanikawa et al., 2002). *Drosophila* mPGES1 is encoded by *mgst1* and is 61% similar at the protein level to its human homolog (UniProt O14684), mPGES2 is encoded by *Su(P)* and is 52% similar (UniProt Q9H7Z7), and cPGES is encoded by *p23* and is 45% similar (UniProt Q15185).

Using available insertional alleles, we assessed the roles of these PGE_2_ synthases in border cell migration. In wild-type follicles, the border cell cluster is in-line with the outer follicle cells throughout S9 (Fig. 1B-B’), indicating on-time migration. Loss of either mPGES1 or mPGES2 does not impact border cell migration (Fig. 1C-D’), whereas loss of cPGES delays border cell migration as the border cell cluster is anterior to the outer follicle cells (Fig. 1E-E’). To quantify border cell migration during S9, we measure the distance of the border cell cluster from the anterior end of the follicle and divide it by the distance of the outer follicle cells; we call this the migration index or MI (Fox et al., 2020; Lamb et al., 2020). On-time migration results in MI of ∼1, whereas delayed migration is <1 (Fig. 1A). Using this method, the MIs in *mPGES1* (1.049, p=0.212) and *mPGES2* (0.958, p=0.529) mutants are similar to wild-type (0.934, Fig. 1F). Conversely, loss of cPGES decreases the MI (Fig. 1F, 0.722, p<0.0001). These data indicate that cPGES-dependent production of PGE_2_ is required for on-time border cell migration.

We next assessed the roles of the synthases in cluster cohesion. Cluster length was not altered by loss of mPGES1, mPGES2 or cPGES (Fig. 1G), suggesting that the cluster elongation phenotype observation when all PG synthesis is lost is not due to the loss of PGE_2_ production.

As cPGES has not been previously studied in *Drosophila*, we next characterized the insertional allele. First, we compared border cell migration in follicles from *cPGES* heterozygotes and homozygotes. Surprisingly, heterozygosity for *cPGES* delays border cell migration (SFig. 1A, MI=0.717, p<0.01). To uncover why the allele has a dominant phenotype, we developed an antibody to cPGES and quantified protein levels by western blot analyses. The allele is a loss of function, as homozygosity for the *cPGES* mutation exhibits 23% of wild-type protein levels, while heterozygosity results in a 30% reduction in protein (SFig. 1B). Together, these data suggest that even mild reductions in cPGES are sufficient to decrease PGE_2_ production enough to impair border cell migration.

### cPGES is required within the substrate for on-time border cell migration

We next asked where cPGES is expressed and localizes to during S9. cPGES is cytoplasmic in all cells of the follicle (SFig. 1C-C’). This staining is reduced in *cPGES* mutant follicles (SFig. 1D-D’). These data indicate that cPGES is present in both the border cells as well as their substrate, suggesting that PGE_2_ could be produced in either or both cell-types.

To determine where PGE_2_ synthesis is required for border cell migration, we used the UAS/GAL4 system to knockdown cPGES by RNAi in either all the somatic cells, including the border cells, or the substrate (Fig. 2A). We first assessed cPGES function in the somatic cells. As expected, the controls (GAL4 only and RNAi only) exhibit on-time migration (Fig. 2B, E; MI = 0.907 and 0.949, respectively). Similarly, somatic knockdown of cPGES does not impact border cell migration (Fig. 2C, E; MI = 0.898). Conversely, knockdown of cPGES in the substrate delays border cell migration (Fig. 2D, E; MI = 0.722, p<0.0001) compared to the controls (GAL4 and RNAi only; MI = 0.940 and 0.949, respectively). To assess knockdown efficiency, we performed immunofluorescence staining for cPGES. Unlike the controls where cPGES is expressed ubiquitously (SFig. 2A), somatic cPGES knockdown retains expression in the substrate but has reduced or absent staining within the somatic cells, including the border cells (SFig. 2B). In the substrate knockdown, cPGES remains expressed in the somatic cells but is reduced in the substrate (SFig. 2C). The migration results were confirmed using the second RNAi line (RNAi-2, SFig. 2D). We also assessed the cell-specific roles of cPGES in regulating cluster morphology. Cluster length is normal when cPGES is knocked down in either the somatic cells or the substrate with either RNAi line (Fig. 2F and SFig. 2E). Together, these data reveal cPGES acts within the substrate to promote on-time border cell migration, but has no role in cluster cohesion.

**Figure 2:**
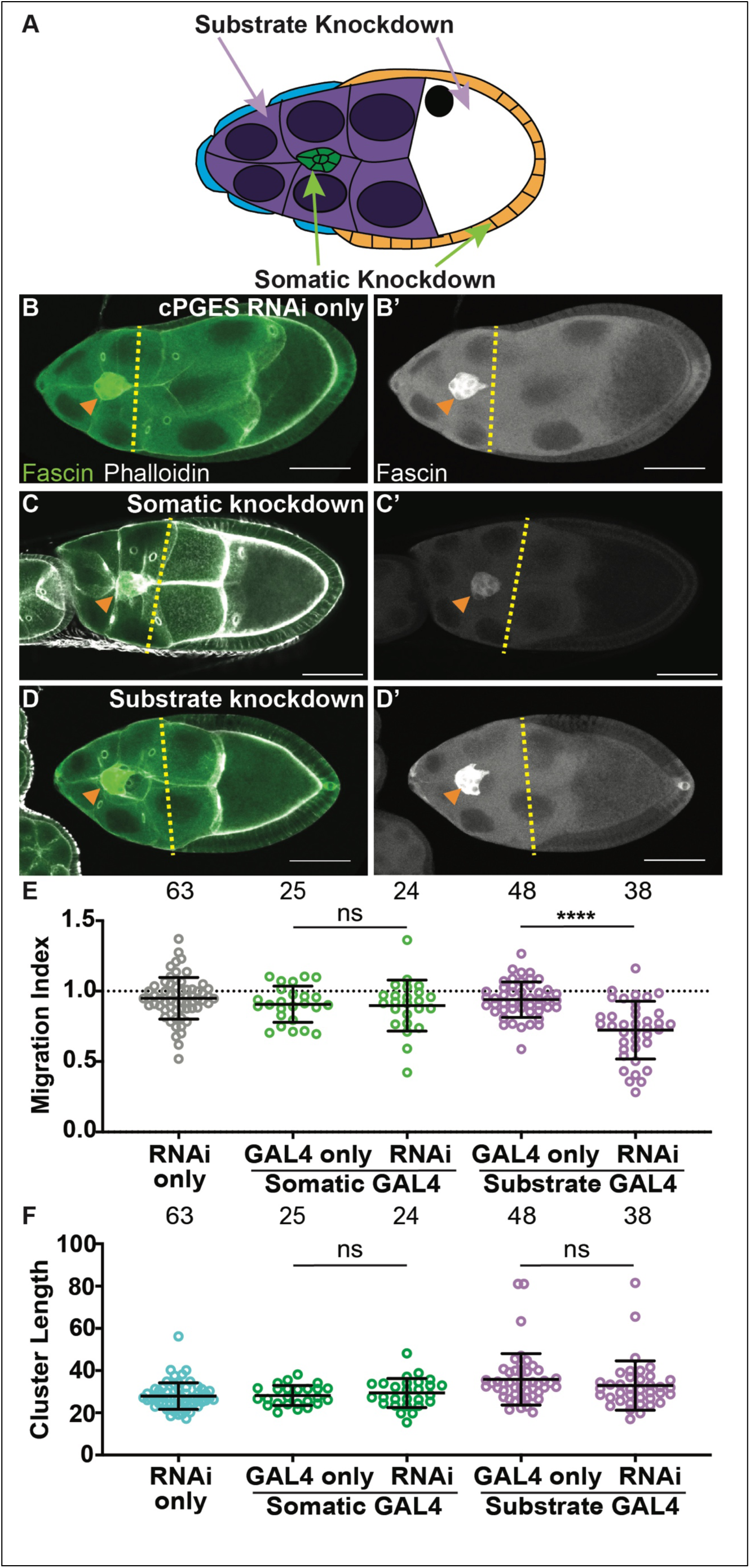
cPGES is required in the substrate for on-time border cell migration. A. Schematic of a S9 follicle indicating the cell-specific knockdown for each GAL4 driver; somatic knockdown occurs in border cells (green) and follicle cells (blue and orange) and substrate knockdown occurs in the nurse cells (purple). B-D’. Maximum projections of 3 confocal slices of S9 follicles stained for Fascin (green in merge) and F-actin (phalloidin, white in merge). Orange arrowheads indicate the border cell cluster and yellow dashed lines indicate the position of the outer follicle cells. Images brightened by 30% to increase clarity. Scale bars = 50μm. B-B’. cPGES RNAi control (*cPGES RNAi/+*). C-C’. Somatic knockdown of cPGES (*c355 GAL4/+; cPGES RNAi/+*). D-D’. Substrate knockdown of cPGES (*osk GAL4/cPGES RNAi*). cPGES RNAi used was HMJ24151. E-F. Graphs of migration index (E) and border cell cluster length (F) for the indicated genotypes. Circle = single follicle; n = number of follicles. In E, the dotted line indicates on-time border cell migration. For E-F, lines = averages and error bars = SD. ns>0.05, **** p<0.0001, unpaired t-test, two-tailed. Like the controls (B-B’, E), somatic knockdown of cPGES exhibits on-time border cell migration (C-C’, E), whereas substrate knockdown delays migration (D-D’, E). Border cell cluster length is unaffected by either somatic or substrate cPGES knockdown (F).

### Akr1B is required for on-time border cell migration

As PG synthesis is required in both the substrate and the migratory cells for migration (Fox et al., 2020), and cPGES acts only in the substrate (Fig. 2 and SFig. 2), a different PG must be produced in the border cells to promote migration. We hypothesized it might be PGF_2α_, as PGF_2α_ drives actin remodeling in later stages of *Drosophila* oogenesis (Tootle and Spradling, 2008; Spracklen et al., 2014), and actin dynamics are critical for border cell migration (Montell, 2003; Montell et al., 2012). In mammals, PGF_2α_ is produced by the aldo-keto reductase proteins Akr1B1, Akr1B10, and Akr1C3 (Banerjee, 2021). In *Drosophila*, the most homologous PGF_2α_ synthase is Akr1B; 85% similar to human Akr1B1 (UniProt P15121). There are two other genes that encode similar aldo-keto reductases: *cg6083*, which appears to be testes-specific, and *cg10638*, which may be a pseudogene that is not expressed (modENCODE). Therefore, we focused on Akr1B.

We used three insertional alleles to assess the role of Akr1B in border cell migration. While wild-type follicles exhibit on-time migration (Fig. 3A, E; MI = 0.948) all three alleles of *akr1B* (PL, d and EY) delay migration (Fig 3B-E; MIs = 0.668, 0.645 and 0.757, respectively, p<0.0001). However, the delay in the *akr1B^EY^* allele was milder, suggesting it is a weaker allele. To test this, we developed an antibody to Akr1B and performed western blot analyses. We find two alleles of *akr1B* (PL and d) reduce protein levels by ∼40%, whereas the *akr1B^EY^* allele only reduces it by ∼9% (SFig. 3A). These data indicate that mild reductions in Akr1B level are sufficient to impair border cell migration. For the rest of the study, we focused on the two stronger alleles. We next assessed the role of Akr1B in regulating cluster morphology. While the *akr1B^PL^* allele has no effect on cluster length, the *akr1B^d0^* allele results in a more compacted cluster (Fig. 3F). Together these findings indicate that Akr1B is required for on-time border cell migration and may play a role cluster morphology.

**Figure 3:**
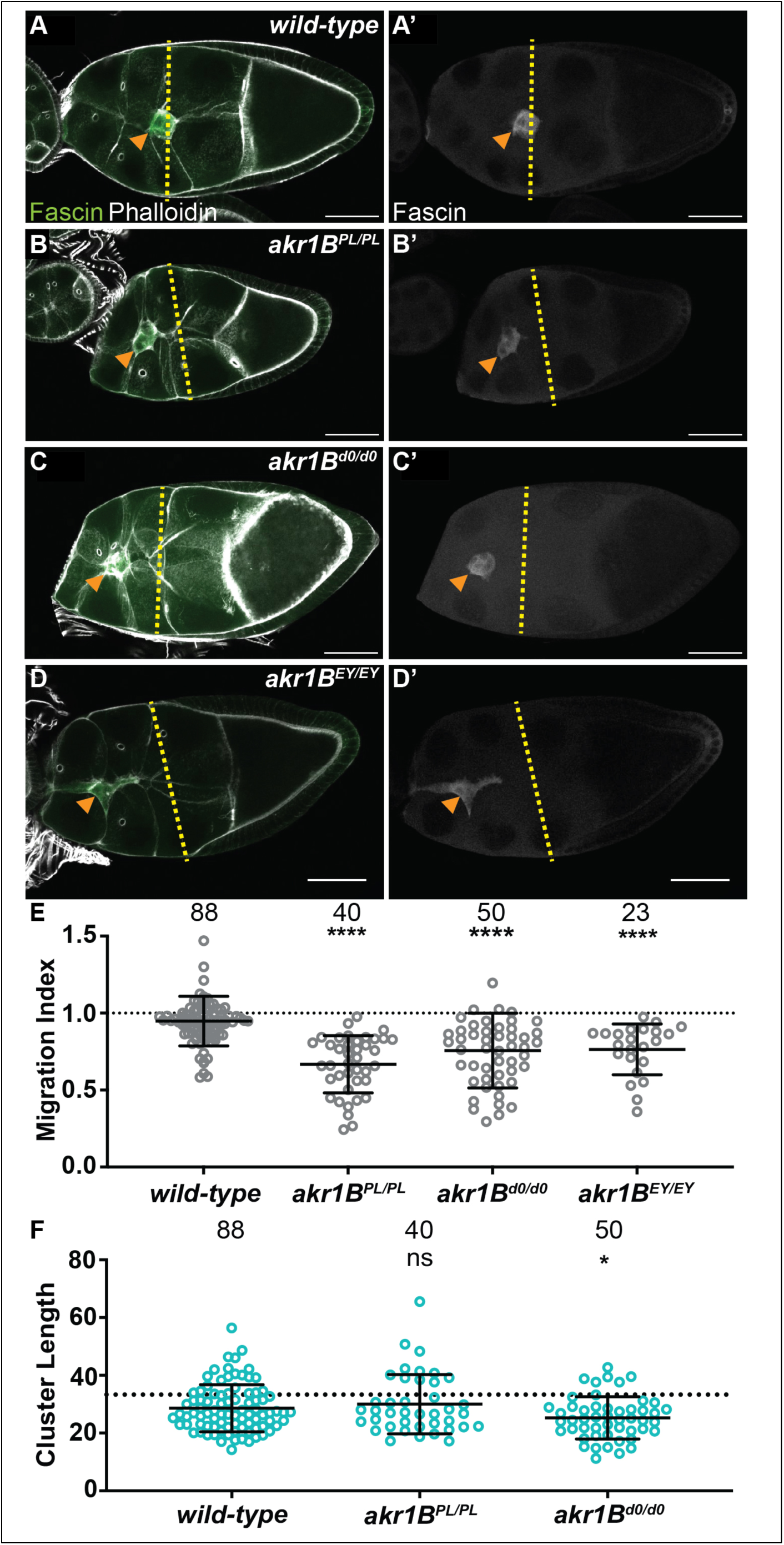
Akr1B is required for border cell migration and cluster morphology. A-D’. Maximum projections of 3 confocal slices of S9 follicles stained for Fascin (green in merge) and F-actin (phalloidin, white in merge). Orange arrowheads indicate the border cell cluster and yellow dashed lines indicate the position of the outer follicle cells. Images brightened by 30% to increase clarity. Scale bars = 50μm. A-A’. *wild-type* (*yw*). B-B’. *akr1B^PL00034/PL00034^*. C-C’. *akr1B^d00405/d00405^*. D-D’. *akr1B^EY07011/EY07011^*. E-F. Graphs of migration index (E) and border cell cluster length (F) for the indicated genotypes; n = number of follicles. In F, the dotted line indicates an on-time border cell migration. For E-F, lines = averages and error bars = SD. ns>0.05, *p<0.05, ** p<0.01 and **** p<0.0001, unpaired t-test, two-tailed. In wild-type S9 follicles, the migrating border cells are in-line with the outer follicle cells (A-A’, E), whereas all three *akr1b* alleles results in delayed migration (B-D’, E). The border cell cluster is more compact in *akr1b^d0/d0^* but not the *akr1b^PL/PL^* allele (F).

### Akr1B is required in the border cells for on-time border cell migration and has cell-specific roles in cluster morphology

To determine where PGF_2α_ synthesis is required for on-time border cell migration, we used the UAS/GAL4 system to knockdown Akr1B in either all the somatic cells, including the border cells, or the substrate. While the controls (GAL4 and RNAi only) exhibit on-time migration (Fig. 4A, D; MIs = 0.930 and 0.936, respectively), RNAi knockdown of Akr1B in the somatic cells slightly delays migration (Fig. 4B, 4D, MI=0.857, p<0.05 and 0.01, respectively). Knockdown of Akr1B within the substrate results in normal migration compared to the two controls (Fig. 4C-D, MI=0.947). These data indicate Akr1B acts in the border cells to promote migration.

**Figure 4:**
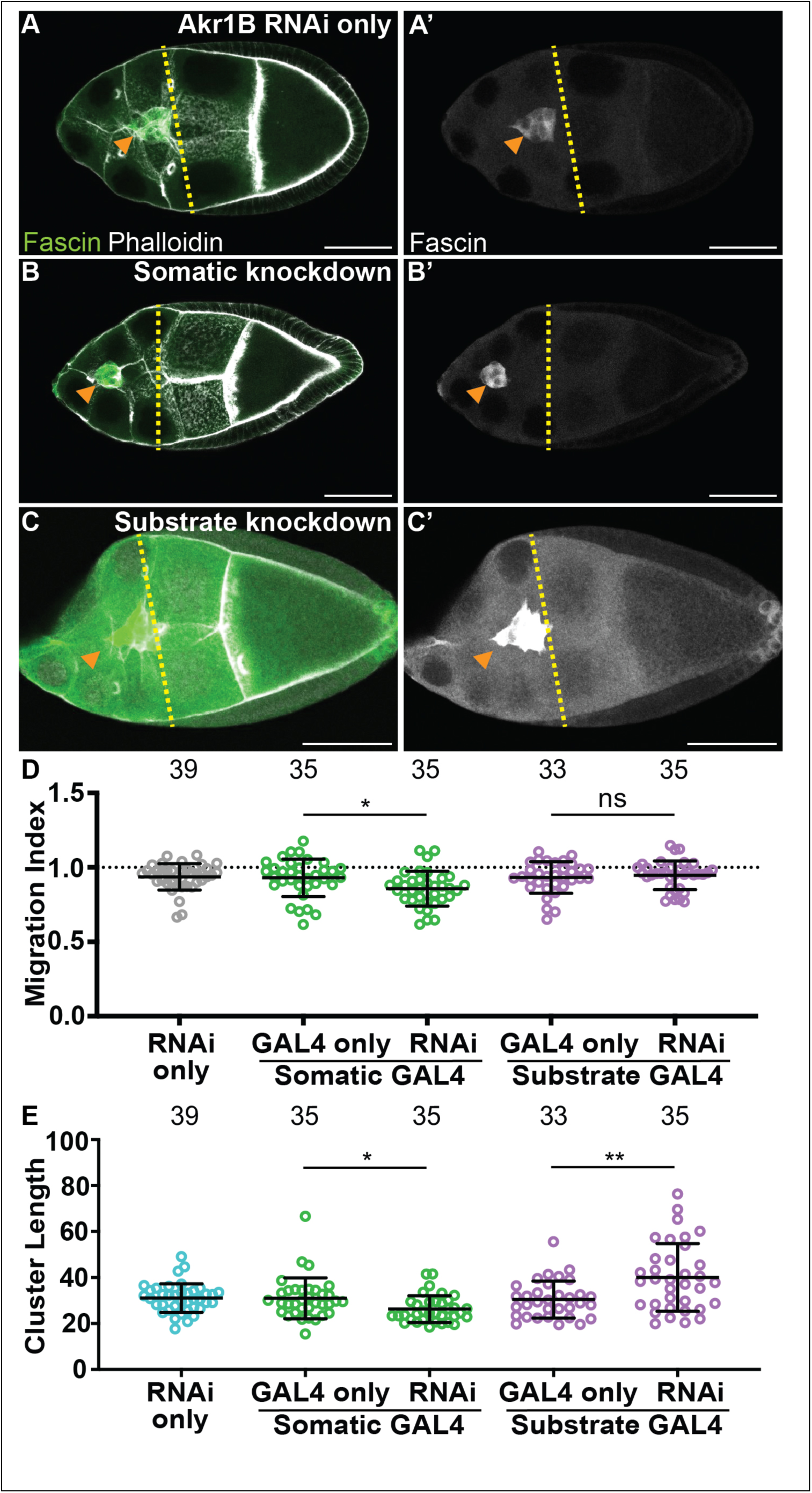
Akr1B is required in the somatic cells for on-time border cell migration, but acts in both the soma and the substrate to regulate cluster morphology. A-C’. Maximum projections of 3 confocal slices of S9 follicles stained for Fascin (green in merge) and F-actin (phalloidin, white in merge). Orange arrowheads indicate the border cell cluster and yellow dashed lines indicate the position of the outer follicle cells. Images brightened by 30% for merge and 85% for Fascin single-channel images to increase clarity. Scale bars = 50μm. A-A’. Akr1B RNAi control (*akr1B RNAi/+*). B-B’. Somatic knockdown of Akr1B (*c355 GAL4/+; akr1B RNAi/+*). C-C’. Substrate knockdown of Akr1B (*osk GAL4/akr1B RNAi*). The *akr1B* RNAi line used was HMS05657. D-E. Graphs of migration index (D) and border cell cluster length (E) for the indicated genotypes; n = number of follicles. In E, the dotted line indicates on-time border cell migration. For D-E, lines = averages and error bars = SD. ns>0.05, * p<0.05, and ** p<0.01, unpaired t-test, two-tailed. Somatic knockdown of Akr1B delays migration (B-B’ compared to A-A’, D), whereas substrate knockdown exhibits on-time migration (C-C’, D). Akr1B somatic knockdown results in more compact clusters but substrate knockdown results in elongated clusters (E).

We next assessed the cell-specific roles of Akr1B in cluster morphology. Somatic knockdown of Akr1B results in a more compact cluster compared to the controls (Fig. 4E, p<0.05 and 0.01, respectively), whereas cluster length is increased in the substrate knockdown (Fig. 4E, p<0.01 and 0.001). These findings are similar to what was observed when dCOX1 was knocked down in the different cell populations (Fox et al., 2020), and therefore, suggests that PGF_2α_ is the PG controlling cluster morphology. Our attempt to confirm the migration and cluster morphology results with a second RNAi line were unsuccessful, as we observed with on-time migration and normal cluster morphology in both somatic and substrate knockdowns (SFig. 4A-B).

### Akr1B, but not cPGES, is required for integrin localization to the border cell membranes

We next sought to identify the mechanisms whereby PGE_2_ and PGF_2α_ promote on-time border cell migration. We first assessed their role in regulating integrins, as integrins are required for border cell migration and cluster cohesion (Dinkins et al., 2008; Menter and Dubois, 2012), and PGs are required for integrin localization to the border cell membranes (Fox et al., 2020). In both wild-type and *cPGES* mutant follicles β-integrin (*Drosophila* Myospheroid) localizes to the membranes (Fig. 5A-A’ and C-C’). Conversely, *akr1b* mutant follicles exhibit a *dCOX1-*like phenotype (Fig. 5B-B’), where integrin staining is diffuse throughout the cytoplasm of the cluster (Fig. 5D-D’). We quantified integrin localization using our previously described method (Fox et al., 2020); see Materials and Methods for details. Loss of *cPGES* is similar to wild-type (p-value=0.632), but *akr1B* mutant follicles have in an integrin intensity ratio below one (Fig. 5E; 0.432, p<0.0001). These results indicate that Akr1B is required for the localization of integrins to the border cell membranes.

**Figure 5:**
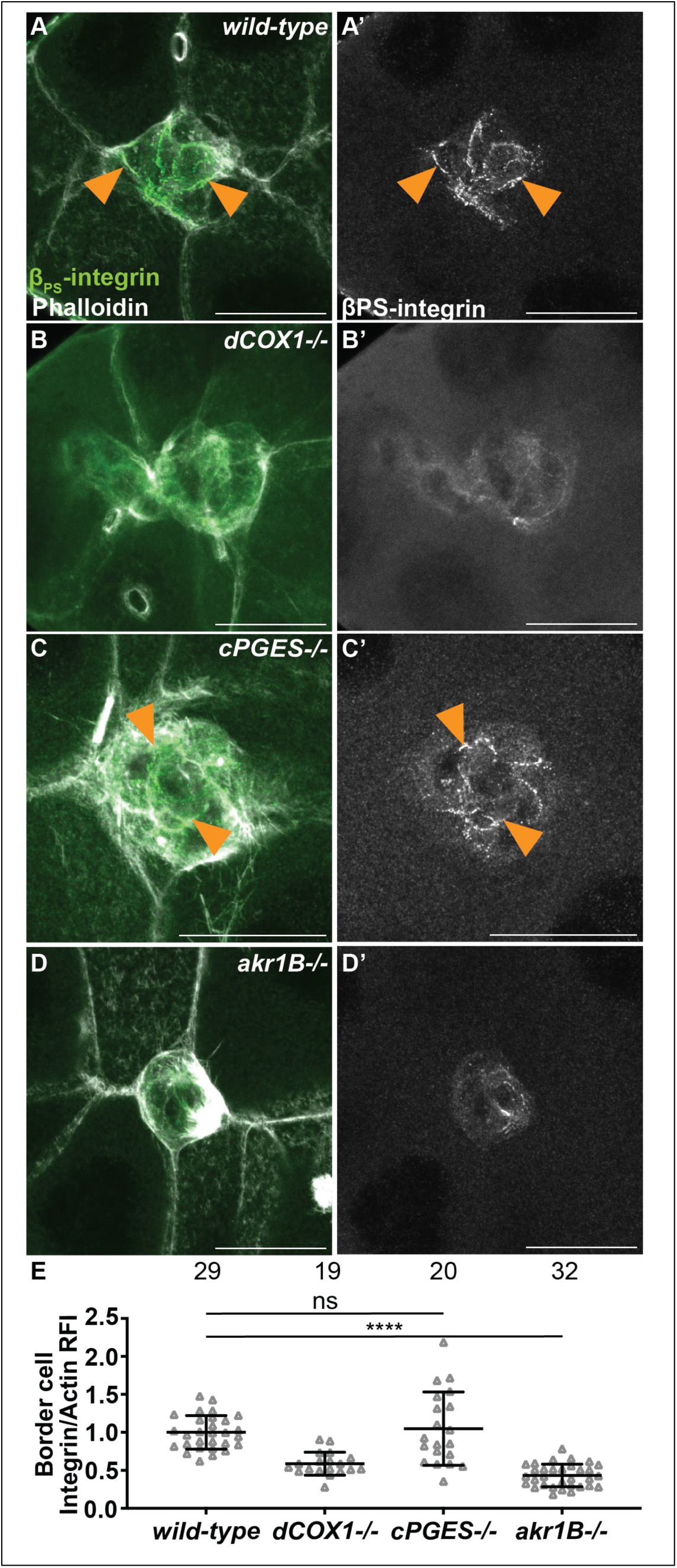
Akr1B is required for integrin localization. A-D’. Maximum projections of 3 confocal slices of S9 follicles stained with β_PS_-integrin (green in merge) and F-actin (phalloidin, white in merge). Arrowheads indicate examples of β_PS_-integrin membrane localization. Images brightened by 30% to increase clarity. Scale bars = 25μm. A-A’. *wild-type* (*yw*). B-B’. *dCOX1-/-* (*dCOX1^f01000/f01000^*). C-C’. *cPGES-/-* (*cPGES^EY05607/^*^EY05607^) D-D’. *akr1B-/-* (*akr1B^d00405/d00405^*). E. Graph of the relative fluorescence intensity (RFI) of β_PS_-integrin to F-actin at the border cell membrane for the indicated genotypes (see Materials and Methods for details); *akr1B-/-* = *akr1B^d00405/d00405^* and *akr1B^PL00034/PL00034^*, and n = number of follicles. Lines = averages and error bars = SD. ns>0.05, **** p<0.0001, unpaired t-test, two-tailed. Like wild-type (A), loss of cPGES (C, E) exhibits integrin localization to the border cell membranes. However, *akr1B* mutants (C, E) have reduced membrane localization that phenocopies *dCOX1-/-* (B, E).

### cPGES limits myosin activity within both the substrate and border cells, whereas Akr1B limits it in only the border cells

The balance of forces between the border cells and their substrate, the nurse cells, is critical for migration, and depends on the level of myosin activity (Majumder et al., 2012; Aranjuez et al., 2016). We previously found that Fascin limits myosin activity within the border cells to control myosin activity in the substrate and thereby, controls substrate stiffness to promote migration (Lamb et al., 2021). As PGs and Fascin act in the same pathway to promote border cell migration (Fox et al., 2020; Lamb et al., 2020), we hypothesize that PG signaling regulates myosin activity. Active myosin is phosphorylated on the myosin regulatory light chain (MRLC) (Vicente-Manzanares et al., 2009; Aguilar-Cuenca et al., 2014). We find that wild-type S9 follicles exhibit a low level of active myosin on the border cell cluster and the substrate (Fig. 6A-A’). We quantified pMRLC relative fluorescence intensity using our previously described method (Lamb et al., 2021); see Materials and Methods for details. Loss of cPGES results in a striking increase in active myosin on both the border cells and their substrate (Fig. 6B-B’, D-E). Whereas in *akr1B* mutants, myosin activity is only increased on the border cells (Fig. 6C-C’, D-E). These findings, together with our cell-specific knockdown results (Figs. 2 and 4), lead to the model that cPGES produces PGE_2_ within the substrate to limit myosin activation and therefore, cellular stiffness, of both the substrate and the migratory cells, whereas Akr1B produces PGF_2α_ to limit the stiffness of only the border cells.

**Figure 6:**
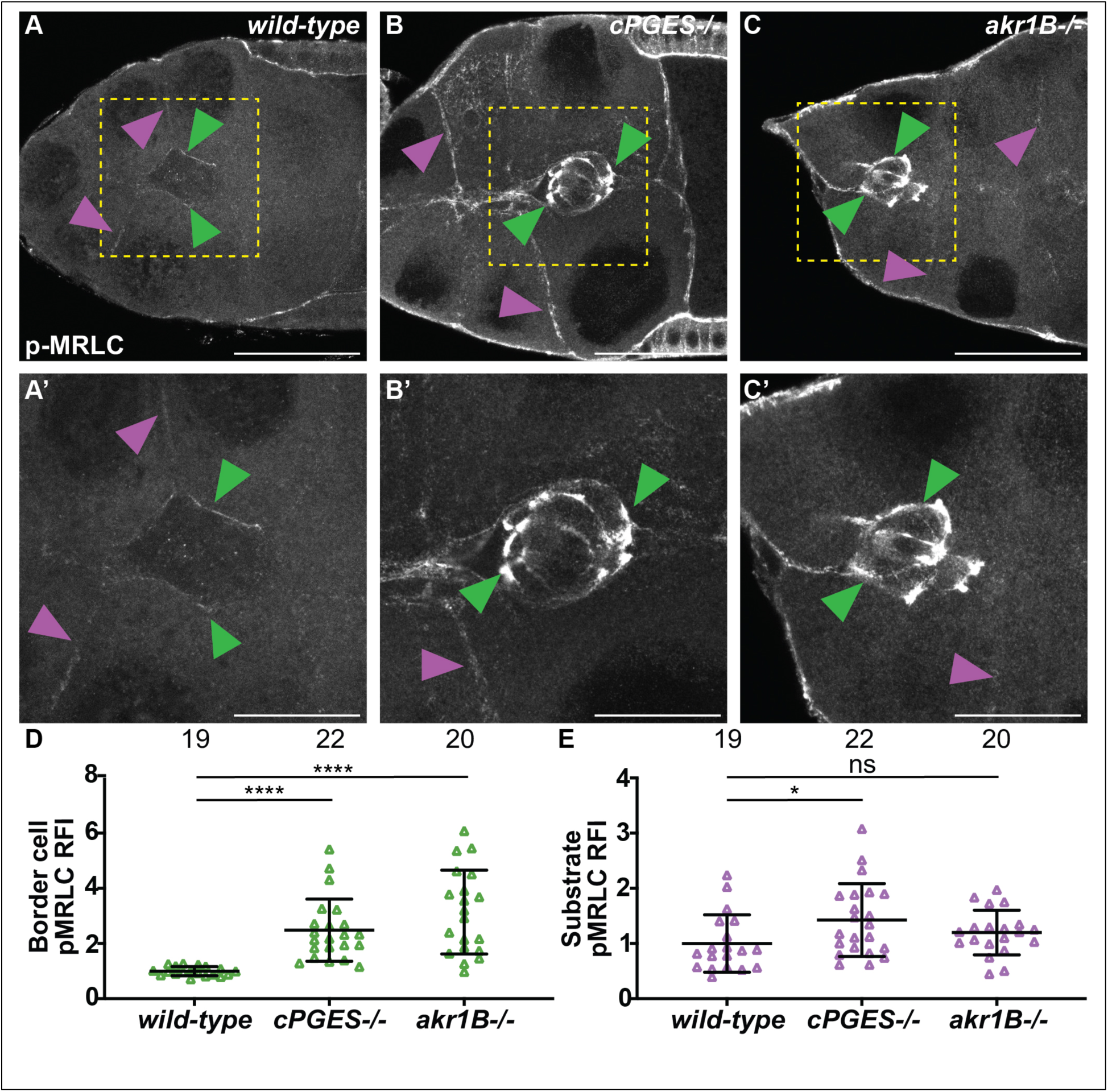
cPGES limits myosin activity within both the border cells and their substrate, whereas Akr1B only limits it within the border cells. A-D’. Maximum projections of 3 confocal slices of S9 follicles stained with pMRLC (white); yellow boxed regions shown at higher resolution in A’-D’. Arrowheads indicate examples of myosin activity on the border cells (green) and substrate (purple). Scale bars = 25μm. Images brightened by 30% to increase clarity. A-A’. *wild-type* (*yw*). B-B’. *cPGES-/-* (*cPGES^EY05607/EY05607^*). C-C’. *akr1B-/-* (*akr1B^PL00034/PL00034^*). D-E. Graphs of relative fluorescence intensity (RFI) of pMRLC on the border cells (D) and the substrate (E) (see Materials and Methods for details); *akr1B-/-* = *akr1B^d00405/d00405^* and *akr1B^PL00034/PL00034^*, and n = number of follicles. Lines = averages and error bars = SD. ns>0.05, **** p<0.0001, unpaired t-test, two-tailed. Loss of cPGES (B-B’) increases myosin activity in both the border cells (D) and the substrate (E) compared to wild-type (A-A’, D-E). Whereas *akr1B* mutants increase myosin activity only in the border cells (C-C’, D-E).

## Discussion

Using *Drosophila* border cell migration as model, we provide the first evidence that both PGE_2_ and PGF_2α_ synthesis, and therefore signaling, are required for a developmental, collective cell migration. We find that the PGE_2_ synthase cPGES is required in the substrate (the nurse cells) but not the border cells for on-time migration (Figs. 1 and 2), whereas PGF_2α_ synthesis by Akr1B is required in the border cells (Figs. 3 and 4). Akr1B acts in both the border cells and the substrate to regulate cluster morphology. Knockdown of Akr1B in the border cells results in compacted clusters, whereas knockdown in the substrate results in cluster elongation (Fig. 4). Potentially contributing to these changes in cluster morphology and to its role in migration, Akr1B promotes integrin-based adhesions on the border cells (Fig 5). Another downstream mechanism whereby both PGs promote cell migration is by limiting myosin activity to control cellular stiffness. Specifically, cPGES limits myosin activity in both the border cells and their microenvironment, while Akr1B limits it only within the border cells. Together these findings reveal that two PGs, PGE_2_ and PGF_2α_, play crucial roles in promoting border cell migration, and that these PGs are produced in distinct locations, the cellular microenvironment and the migrating cells, respectively (Fig. 7).

**Figure 7:**
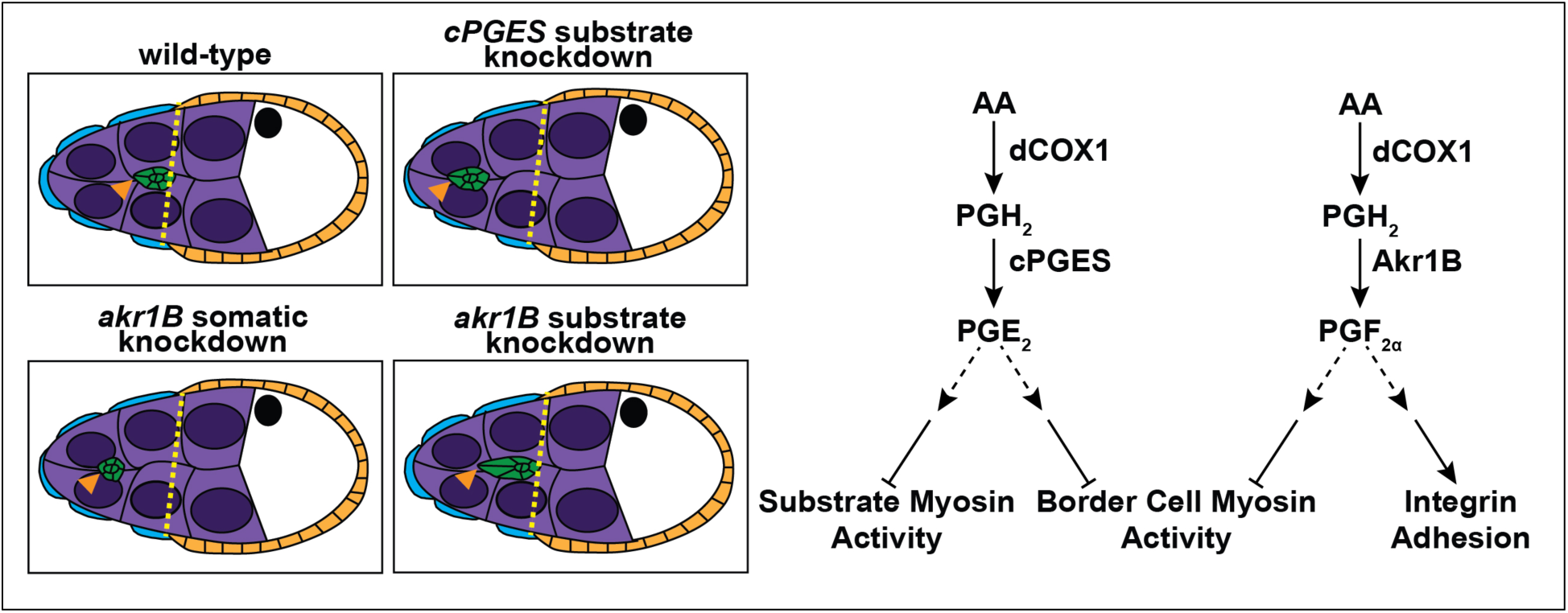
Schematics of the roles of PGE_2_ and PGF_2α_ in border cell migration. Schematics of S9 follicles of the indicated genotypes (left) and pathway diagrams (right). In *wild-type* follicles the border cells (green) are in-line with the position of the outer follicle cells (orange and yellow dashed line). Substrate knockdown of *cPGES* and somatic knockdown *akr1B* delay migration. Thus, cPGES is required in the substrate and Akr1B is required in the somatic cells for on-time migration. Additionally, Akr1B regulates border cell cluster cohesion, as somatic knockdown results in more compact clusters while substrate knockdown results in elongated clusters. Our data supports the model that cPGES in the substrate regulates border cell migration by limiting myosin activity in both the substrate and the border cells, whereas Akr1B only limits myosin activity in the border cells. Akr1B also promotes integrin-based adhesions on the border cells. Which cells produce Akr1B to regulate these different downstream effectors remains to be determine.

### Mild reductions in PG synthase levels impair migration

Incomplete loss of either cPGES or Akr1B delays border cell migration. Migration is delayed by heterozygosity for *cPGES*, which reduces protein levels by 30% (Fig. 1 and SFig. 1B), and weak alleles of *Akr1B*, which reduce protein levels by 9-40% (Fig. 3 and SFig. 3A). These findings suggest that mild decreases in PG-type specific synthase levels have striking impacts on PG production and downstream PG signaling. Supporting this idea, siRNA knockdown of cPGES in colon cancer cells retains 36% of the protein level but impairs invasiveness (Cano et al., 2015). Similarly, incomplete knockdown of mPGES1 impairs carcinoma cell migration (Kamei et al., 2009; Lu et al., 2012; Jongthawin et al., 2014). Studies in breast cancer cells support that siRNA knockdown of Akr1B1 retains some protein expression but results in an almost complete loss of PGF_2α_ production and impairs migration and invasion (Wu et al., 2017). These data lead us to speculate that heterozygosity for *cPGES* and mild reductions in Akr1B reduce PGE_2_ and PGF_2α_ levels, respectively, below that needed to promote border cell migration. Directly testing this idea will require the development of a method that allows cell-specific levels of individual PGs to be assessed within S9 follicles, as current methods of quantifying PGs – enzyme-linked immunosorbent assay or high-performance liquid chromatography-mass spectrometry – require large amounts of cells and do not provide cellular resolution.

### PGE_2_ produced in the microenvironment promotes cell migration

Numerous studies support that PGE_2_ promotes migration. However, it remains largely unclear which cells produce PGE_2_ versus which cells respond to the PGE_2_ signal. Indeed, studies in zebrafish reveal that COX activity and PGE_2_ synthesis/signaling are required for gastrulation (Cha et al., 2005; Cha et al., 2006). However, the cell-specific roles are unknown as the studies employed COX inhibitors and whole organism knockout of COX1 or a PGE_2_ synthase.

Studies measuring PGE_2_ production from tumors and cancer cell lines led to the idea that PGE_2_ is produced and signals autocrinely within cancer cells to drive migration. For example, high COX2 activity is associated with breast cancer metastasis, and breast cancer cell lines with high COX2 levels and PGE_2_ production exhibit increased migration and invasion that are impaired by COX2 inhibition (Singh et al., 2005). Further, studies reveal exogenous PGE_2_ promotes the migration of lung, colon, and prostate cancer cells (Kim et al., 2010; Fujino et al., 2011; Vo et al., 2013).

However, there is growing evidence that cells within the cancer microenvironment produce PGE_2_ to promote cancer migration and invasion (Elwakeel et al., 2019). Use of a co-culture system revealed that colon cancer cells secrete interleukin-1 to activate PGE_2_ production in mesenchymal stem cells (MSCs). This PGE_2_ signals both autocrinely within the MSCs and paracrinely to the cancer cells to drive dedifferentiation and invasion (Li et al., 2012). Similarly, in head and neck squamous cell carcinoma, fibroblasts upregulate PGE_2_ production which then induces tumor cell proliferation and migration (Alcolea et al., 2012). Genetic deletion of mPGES1 in mice suppresses two models of colon cancer, however, mPGES1 is weakly expressed in the tumor, but highly expressed in the adjacent colonic mucosa, raising the possibility that PGE_2_ is produced within the microenvironment (Nakanishi et al., 2008; Nakanishi et al., 2011). Further, mPGES1 knockdown lung cancer xenografts grow slower, but this is enhanced by implantation into mPGES1 knockout mice; markers of invasion are also decreased (Kamei et al., 2009). These studies support that PGE_2_ produced in the microenvironment contributes to cancer progression, migration, and invasion.

PGE_2_ also acts paracrinely to promote immune cell migration. In a breast cancer xenograft model, overexpression of COX2 induces PGE_2_ production which recruits regulatory T-cells (Karavitis et al., 2012). Exogenous PGE_2_ is required for dendritic cells to respond to chemokines and chemoattractants (Legler et al., 2006), and reorganizes the actin cytoskeleton to promote migration (Diao et al., 2021). Further, there is evidence that a gradient of PGE_2_ is critical for regulating the state of macrophages, from driving migration at low levels to promoting phagocytosis at high levels (Osma-Garcia et al., 2016).

Given these studies, it is not surprising that we find that PGE_2_ synthesis is required in the microenvironment for border cell migration (Fig. 2). Whether all of the nurse cells produce PGE_2_ or produce the same level of PGE_2_ remains unknown, but it is tempting to speculate there may be a gradient from low to high PGE_2_ levels along the anterior to posterior axis that promotes border cell migration.

Our finding that cPGES is required in the microenvironment for on-time border cell migration, initially contradicts our finding that when all PG synthesis is blocked by RNAi knockdown of dCOX1 in the microenvironment, border cell migration was on-time. This knockdown of dCOX1 was achieved by using an RNAi line that is normally unable to be expressed in the germline, but can be expressed when combined with reduced Hsp70 (DeLuca and Spradling, 2018). This method likely resulted in a weak knockdown of dCOX1 and we speculate this level of knockdown was not sufficient to limit PG production enough to uncover the microenvironment roles in promoting migration.

We find that cPGES, but not mPGES1 or mPGES2, promotes border cell migration. This finding was unexpected, as mPGES1 is widely implicated in cell migration during both development (Cha et al., 2005; Cha et al., 2006) and cancer (Nakanishi et al., 2008; Kamei et al., 2009; Nakanishi et al., 2011). However, cPGES is also implicated in cancer migration (Cano et al., 2015). One possible reason for our results is that the mPGES1 and mPGES2 alleles do not reduce protein levels sufficiently to cause a phenotype. We think this is unlikely given that heterozygosity for *cPGES* is sufficient to delay migration (SFig 1A). Therefore, we favor the model that cPGES is the primary synthase responsible for producing PGE_2_ within the microenvironment to promote border cell migration.

### PGF_2α_ produced in the migratory cells promotes migration

PGF_2α_ is an understudied PG but has been implicated in acting both autocrinely and paracrinely to promote migration. In breast cancer cell lines, knockdown or inhibition of Akr1B1 decreases, whereas overexpression increases migration and invasion (Wu et al., 2017). In an endometrial cancer cell line, exogenous PGF_2α_ or PGF_2α_ receptor agonist increases migration (Sales et al., 2008). Further, in patients with colon cancer, high expression of Akr1B1 is associated with enhanced motility and poor clinical outcome (Demirkol Canli et al., 2020). These data suggest that PGF_2α_ produced in both the cancer cells and their microenvironment contributes to migration. However, the literature suggests PGF_2α_ acts paracrinely to regulate immune cell migration. PGF_2α_ produced by endothelial cells in the contexts of hypoxia (Arnould et al., 2001) or by endometrial cancer cells promotes neutrophil migration (Wallace et al., 2009). We find that during *Drosophila* border cell migration Akr1B is required within the migratory cells for on-time migration (Fig. 4). One caveat to our findings is that the available RNAi lines may produce only weak knockdown. Therefore, Akr1B may act in the microenvironment to regulate border cell migration, but to observe such a role we need a stronger loss of Akr1B than we can currently achieve. However, we think this is unlikely as the *akr1b^EY^*allele only results in a 9% reduction in protein but delays migration (Fig. 2 and SFig. 3).

### PGF_2α_ regulates cluster cohesion

In addition to promoting on-time border cell migration, PGs also regulate cluster cohesion. Loss of dCOX1 results in cluster elongation, with clusters sometimes breaking apart (Fox et al., 2020). Similar cluster elongation is seen when dCOX1 is knocked down in the substrate. However, knockdown in the border cells results in cluster compaction, revealing PGs have cell-specific roles in controlling cluster cohesion. Here we find that the PG responsible for regulating cluster cohesion is PGF_2α_. Global reduction of the PGF_2α_ synthase Akr1B results in cluster compaction, as does knockdown in the border cells (Figs. 3 and 4). However, knockdown in the substrate causes cluster elongation (Fig. 4). Together these data, along with our findings on dCOX1 (Fox et al., 2020), lead to the model that PGF_2α_ produced within the border cell cluster acts to limit cluster cohesion, whereas PGF_2α_ produced in the substrate promotes cluster cohesion.

While it remains unclear how PGF_2α_ signaling regulates cluster cohesion, our data suggest a few possible mechanisms. First, *akr1b* mutants reduce integrin-based adhesions on the border cells (Fig. 5). RNAi knockdown of either subunit of the integrin receptor delays border cell migration, and, when combined with reduced JNK signaling, elongates clusters (Dinkins et al., 2008; Llense and Martin-Blanco, 2008). Therefore, one means by which PGF_2α_ signaling may control cluster cohesion is by tightly regulating integrin-based adhesions. Supporting this idea, in cancer, PGs promote integrin adhesion stability (Mayoral et al., 2005; Bai et al., 2009; Liu et al., 2010). Second, these morphology changes may be due to PGF_2α_ signaling controlling actin cytoskeletal remodeling within the border cells. Indeed, during later stages of *Drosophila* oogenesis PGF_2α_ maintains cortical actin integrity and promotes actin bundle formation (Tootle and Spradling, 2008; Groen et al., 2012; Spracklen et al., 2014). Third, either by regulating the actin cytoskeleton or by other means, PGF_2α_ may control cluster cohesion by modulating cellular stiffness. We find *akr1b* mutants exhibit increased border cell stiffness, as seen by increased myosin activity (Fig. 6). The balance of forces between the border cells and their substrate must be tightly regulated for normal cluster morphology, as misbalanced forces cause cluster elongation (Majumder et al., 2012; Cai et al., 2014; Aranjuez et al., 2016). PGF_2α_ regulation of both cellular stiffness and actin cytoskeletal dynamics may modulate integrin-based adhesions. Thus, all three mechanisms may contribute to PGF_2α_ control of cluster cohesion.

### PGs regulate the balance of forces to promote migration

Cell migration depends on both the stiffness of the migratory cells and their microenvironment, and the balance of those forces (Kai et al., 2016). Numerous studies have shown that substrate stiffness regulates migratory cell stiffness and ability to migrate (Aguilar-Cuenca et al., 2014; Barriga et al., 2018); this is particularly evident in cancer migration and metastasis (Gasparski et al., 2017; Oakes, 2018; Eble and Niland, 2019; Ren et al., 2021). Evidence is also emerging that migrating cells influence their microenvironment. For example, migrating cells degrade extracellular matrix (ECM) to promote migration (Wolf et al., 2007); this likely decreases microenvironment stiffness. Migrating cells can also increase the stiffness of the microenvironment, by pulling on and aligning ECM fibers (Hall et al., 2016; van Helvert and Friedl, 2016). Further, cancer cells induce changes in the stroma, including increasing fibrosis and, thereby, stiffening the tissue (van Helvert et al., 2018; Chandler et al., 2019; Piersma et al., 2020). Ultimately, this increase in the stiffness of the microenvironment promotes cell migration, increasing the force generation in the migratory cells by a process termed mechanoreciprocity (Cox and Erler, 2014; van Helvert et al., 2018). Such a coordinated and interdependent balance of forces is seen between the border cells and their microenvironment, the nurse cells (Majumder et al., 2012; Aranjuez et al., 2016; Lamb et al., 2021).

A key regulator of cellular stiffness is myosin, a force generating actin motor (Vicente-Manzanares et al., 2009; Aguilar-Cuenca et al., 2014). Indeed, myosin controls the stiffness of both migrating cells and their cellular substrates (Lo et al., 2000; Vicente-Manzanares et al., 2009; Mohan et al., 2015). Further, myosin serves as a force sensor, driving the cellular response to applied forces (Butcher et al., 2009; Vicente-Manzanares et al., 2009; Aguilar-Cuenca et al., 2014). Myosin plays these important roles during border cell migration (Majumder et al., 2012; Aranjuez et al., 2016; Lamb et al., 2021). When myosin activity is severely increased in the nurse cells, the microenvironment, it increases active myosin in the border cells and delays migration (Aranjuez et al., 2016). Increasing myosin activity on the border cells also drives myosin activation and stiffening of the nurse cells. This migratory cell influence on the microenvironment depends of Fascin (Lamb et al., 2021). This function of Fascin is likely regulated by PG signaling, as Fascin is a downstream effector of PGs during border cell migration (Fox et al., 2020).

We find that PGE_2_ and PGF_2α_ synthesis have cell-specific roles in regulating myosin activity (Fig. 6). Loss of cPGES results in increased myosin activation on both the border cells and the nurse cells, whereas reduction in Akr1B only increases it on the border cells. This loss of mechanoreciprocity could be due to insufficient reduction in Akr1B and thereby, PGF_2α_ levels. Alternatively, it could indicate cell-specific roles of the different PGs. Taking our cell-specific knockdown findings into account (Figs. 2 and 4), we speculate: Akr1B-dependent PGF_2α_ production within the border cells limits myosin activity and border cell stiffness. PGF_2α_ synthesis may also required for the nurse cells to appropriately respond to the forces placed on them. PGE_2_ synthesis by cPGES in the nurse cells signals to the border cells to modulate border cell stiffness, which in turn controls nurse cell stiffness. We hypothesize that both PGE_2_- and PGF_2α_-dependent regulation of myosin activity occurs, at least in part, via modulating Fascin activity.

The role of PGs in regulating myosin activity is likely conserved. In colonic lamina propria fibroblasts, PGE_2_ signaling is required for reducing myosin activity to allow cell polarization and migration during wound healing (Rieder et al., 2010). PGE_2_ regulates myosin activation in dendritic cells, controlling their maturation (van Helden et al., 2008). PGF_2α_ promotes myosin activation in muscle cells, driving their contraction (Ansari et al., 2004; Xu et al., 2015); how it influences myosin activity in other cells remains unknown. Future studies on *in vivo* migrating cells, like the border cells, are needed to uncover the roles of distinct PGs in modulating myosin activity and cellular stiffness to promote migration.

### Do PGE_2_ and PGF_2α_ signal at different times during border cell migration?

Our data shows that both PGE_2_ and PGF_2α_ synthesis are required for on-time border cell migration. However, it remains unknown whether they signal simultaneously or at distinct times, and whether one PG induces the production of the other. Supporting the latter possibilities, in colorectal tumor cells PGF_2α_ signaling induces the production of PGE_2_ (Stamatakis et al., 2015). If this occurs during border cell migration, it could help explain why small reductions in Akr1B levels result in such striking delays in border cell migration (Fig. 3 and SFig. 3). However, if this were the only mechanism controlling PGE_2_ production, one would predict loss of Akr1B would phenocopy loss of cPGES and exhibit increased myosin activity in both the border cells and nurse cells (Fig. 6). It is also possible that force transmission from the border cells to the nurse cells, which likely occurs by both PGF_2α_-dependent and independent mechanisms, activates PGE_2_ production. Indeed, cytoplasmic phospholipase A2 (cPLA2) is activated by mechanical signaling, resulting in the release of arachidonic acid, the substrate for all PG production (Enyedi et al., 2016; Lomakin et al., 2020); substrate release is the rate limiting step in PG synthesis (Funk, 2001; Tootle, 2013). Future studies, in conjunction with developing methods for visualizing the timing of PG synthesis and signaling, are needed to determine the interplay between PGE_2_ and PGF_2α_ in border cell migration.

## Conclusion

The field’s understanding of cell-specific roles of individual PGs in cell migration has been limited by the widespread use of ubiquitously perturbing PG synthesis and signaling components, and by studying cellular responses to exogenously supplied PGs. *Drosophila* border cell migration provides an *in vivo*, physiological system to decipher the cell-specific roles of different PGs in promoting collective cell migration, allowing the separation of roles within the migratory cells versus their microenvironment (Fig. 7). Here we find that the most widely studied PG, PGE_2_, is not synthesized by the migratory cells, but is produced by cPGES in the microenvironment to promote border cell migration. Further, migration requires PGF_2α_, an understudied PG, to be produced by Akr1B in the border cells. These findings call for a reassessment of the cellular site of PGE_2_ activity, and for widespread examination of the roles of PGF_2α_ in cell migration, from development to cancer metastasis. Our work suggests both PGs promote migration by controlling myosin activity and cellular stiffness, but whether they do so by the same or different mechanisms remains unknown. Further, we find PGF_2α_, but not PGE_2_ synthesis, is required for integrin-based adhesions. It will be important to determine whether these downstream mechanisms of PGE_2_ and PGF_2α_ signaling are conserved across some or all collective cell migrations.

## Acknowledgements

We thank Omar Rabab’h for efforts during his lab rotation, the Dunnwald lab for helpful discussions, and the Tootle lab for helpful discussions and careful review of the manuscript. Stocks obtained from the Bloomington Drosophila Stock Center (NIH P40OD018537) were used in this study. Resources provided by FlyBase (NIH U41HG-000739) were used for this study. At the University of Iowa, Information Technology Services – Research Services provided data storage support.

## Materials and Methods

### Reagents and resources

See Table S1 for detailed information on the reagents used in these studies and Table S2 for the specific genotypes used in each figure panel. All raw data used in this study can be found in Table S3.

### Fly stocks

Fly stocks were maintained on cornmeal/agar/yeast food at 21°C, except where noted. Before immunofluorescence staining, newly eclosed flies were fed wet yeast paste every day for 2-4 days. Unless specified, *yw* (BDSC 1495) was used as the control. The following stocks were obtained from the Bloomington *Drosophila* Stock Center: *c355 GAL4* (BDSC 3750), *c306 GAL4* (BDSC 3743), *mgst1*^KG04713^ (BDSC 13839), *Su(P)^EY13245^* (BDSC 20866)*, p23^EY05607^*(BDSC 16661)*, akr1B^PL00034^* (BDSC 19594)*, akr1B^EY07011^* (BDSC 16777), *p23 RNAi* HMJ24151 (BDSC 62911), *p23 RNAi-2* GL01292 (BDSC 41862), *akr1B* RNAi HMS05657 (BDSC 67838), *akr1B RNAi-2* HMC05226 (BDSC 62219) and *UAS Dicer 2* (BDSC 24651). The following stocks were obtained from the Exelexis Stock Center: *mgst1^d10243^*, *akr1B^d00405^* and *pxt^f01000^* (Thibault et al., 2004). The *oskar GAL4* line (second chromosome; BDSC 44241) was a generous gift from Anne Ephrussi (European Molecular Biology Laboratory; (Telley et al., 2012)). Expression of the RNAi lines were achieved by crossing to *c355 GAL4* or *oskar GAL4*, maintaining fly crosses at 21°C and maintaining progeny at 29°C for 5-6 days. *UAS Dicer* was used in combination with *c355* to enhance RNAi efficiency were noted in the figure legends.

### Immunofluorescence

*Drosophila* ovaries (5-8 pairs per sample) were dissected into room temperature Grace’s insect medium (Lonza). Ovaries were fixed for 10 min using 4% paraformaldehyde diluted in Grace’s medium. Samples were washed six times for 10 min each at room temperature in antibody wash (1X phosphate-buffered saline [PBS], 0.1% Triton X and 0.1% bovine serum albumin [BSA]). Primary antibodies were diluted in antibody wash and incubated overnight at 4°C, except for β_PS_-integrin which was incubated for ∼48-72 hours at 4°C. The following monoclonal antibodies were obtained from the Developmental Studies Hybridoma Bank (DSHB), created by the NICHD of the NIH and maintained at The University of Iowa, Department of Biology, Iowa City, IA : mouse anti-Fascin 1:50 (sn7c, Cooley, L; AB_528239; (Kelly Cant, 1994)) and mouse anti-β_PS_-integrin 1:10 (CF.6G11, Brower, D; AB_528310; (Danny L. Brower, 1984)). A rabbit polyclonal antibody (Genscript) produced against full-length *Drosophila* p23 (Q9VH95) was used at 1:1000. After six washes in antibody wash (10 min each), samples were incubated in secondary antibodies overnight at 4°C. The following secondaries were used at 1:500: AF488∷goat anti-mouse (AB_2534069), AF568∷ goat anti-mouse (AB_2534072), AF488 goat anti-rabbit (AB_2576217), and AF568∷ goat anti-rabbit (AB_ 2534102) (Thermo Fischer Scientific). Alexa Fluor 586- or Alexa Fluor 647-conjugated phalloidin (A12380 and A22287; Thermo Fischer Scientific) diluted 1:250 were included in both primary and secondary antibody incubations. Following six washes in antibody wash (10 min each), 4’,6-diamidino-2-phenylidole (DAPI; 5 mg/mL; D3571; Thermo Fischer Scientific) staining was performed at a concentration of 1:5000 in 1X PBS for 10 min at room temperature. Samples were then rinsed in 1X PBS and mounted on slides in 1 mg/mL phenylenediamine in 50% glycerol, pH 9 (Platt and Michael, 1983). All experiments were performed a minimum of three independent times.

Phospho-myosin regulator light chain (pMRLC) staining was performed using a protocol provided by the McDonald Lab (Majumder et al., 2012; Aranjuez et al., 2016). Briefly, ovaries were fixed for 20 min at room temperature in 8% paraformaldehyde in 1x PBS and 0.5% Triton X-100. Samples were blocked for 30 min at room temperature in Triton antibody wash (1X PBS, 0.5% Triton X-100, and 5% BSA). Primary antibodies, rabbit anti-pMRLC (S19, 1:100; AB_330248; Cell Signaling), mouse anti-Hts (1:50; Lipshitz, H; DSHB; AB_ 528070; (Zaccai and Lipshitz, 1996)) and mouse anti-FasIII (1:50, Goodman,C; DSHB; AB_ 528238; (Patel et al., 1987)), were diluted in Triton antibody wash and incubated for ∼48-72 hours at 4°C. Following six washes in Triton antibody wash (10 min each), secondary antibody staining, washes, and DAPI staining was performed and samples were mounted as described above.

### Image acquisition and processing

Microscope images for fixed and stained *Drosophila* follicles were taken using LAS SPE Core software on a Leica TCS SPE mounted on a Leica DM2500 using ACS APO 20x/0.6 IMM Corr -/D objective (Leica Microsystems), Zen Software on Zeiss 700 LSM mounted on an Axio Observer.Z1 using a Plan-Apochromat 20x/0.8 working distance= 0.55 M27 or EC-Plan_Neo_Fluar 40x/1.3 oil objective, or Zeiss 880 mounted on Zeiss Axio Observer.Z1 using Plan-Apochromat 20x/0.8 working distance= 0.55 M27, Plan-Apochromat 40x/1.3 oil DIC working distance= 2.0 (SCR_013672) (Carl Zeiss Microscopy). S9 follicles were identified by their size (∼150μm-250μm) and morphology, including, the location of the outer follicle cells and the border cell cluster. The beginning of S10 was defined as when the anterior most outer follicle cells reached the nurse cell-oocyte boundary and flattened. Maximum projections, merge, rotation, and cropping were performed using ImageJ software (FIJI, RRID: SCR 002285, (Abramoff et al., 2004)). All images shown were brightened by 30% in Photoshop (Adobe, RRID: SCR 014199), except where noted, to improve visualization and figures were made using Illustrator (Adobe, RRID: SCR 010279).

### Quantification of border cell migration and cluster length

Quantification of the MI was performed as described previously (Fox et al., 2020; Lamb et al., 2020). Briefly, measurements of S9 follicles were performed using ImageJ software (Abramoff et al., 2004) on maximum projections of 2-4 confocal slices of follicles stained for Fascin and phalloidin. A line segment was used to measure the distance in microns from the anterior end of the follicle to the leading edge of the border cell cluster; this was defined as the border cell distance. Another line segment was used to measure the distance from the anterior end of the follicle to the anterior end of the outer follicle cells: this was defined as the outer follicle cell distance. The entire follicle length was also measured along the anterior-posterior axis. The migration index was calculated by dividing the border cell distance by the follicle cell distance. Cluster length was determined by measuring the distance from the front to the rear of the border cell cluster (detached cells were not included). Data were compiled in and calculations were performed in Excel (Microsoft, RRID: SCR 016137), and graphs were generated and statistical analyses performed using Prism (GraphPad Software, RRID SCR 002798).

### Quantification of Integrin Localization

Integrin analysis was performed using the method described previously (Fox et al., 2020). Briefly, integrin intensity was measured from single confocal slices of immunofluorescence images of fixed *Drosophila* follicles. The “straight line” function was used to draw lines between 4-7 microns at three different locations on the border cell membranes, and highest fluorescence intensity value was measured for β_PS_-integrin and phalloidin. The integrin value was divided by the phalloidin value, the average was calculated for the three segments, and then the average was normalized to the overall *wild-type* average. Data were compiled in and calculations were performed in Excel, and graphs were generated and statistical analyses performed using Prism.

### pMRLC quantifications

pMRLC analysis was performed as previously described (Lamb et al., 2021). Briefly, intensity measurements were performed on maximum projections of 3 confocal slices of 40x confocal images at a 2x zoom using ImageJ software ((Abramoff et al., 2004). The fluorescence intensity of pMRLC on the border cells was measured by tracing the shape of the border cell cluster and obtaining the highest intensity value of the pMRLC staining. The same shape was used to measure the background within the substrate, which was substracted. pMRLC values were normalized to the *wild-type* average. To determine the substrate pMRLC intensity, three-line segments (14-16μm) at different locations within the follicle were used to measure both pMRLC and phalloidin fluorescent intensity peaks. Values were normalized by division with the phalloidin intensities and averaged for one sample. Averages were then normalized to the *wild-type* average. Data was compiled in and calculations were performed in Excel, and graphs were generated and statistical analyses performed using Prism.

### Western Blot

Whole ovary pairs (5 total per sample) were dissected at room temperature in 1xPBS and transferred to a 1.5mL tube containing 80μl 1xPBS. 20μl 5x Laemmli buffer was added and lysis was performed by grinding tissue with plastic pestles (RNase-free disposable pellet pestles; Thermo Scientific). Samples were boiled for 10 min and briefly spun down before loading. Western Blots were performed using standard methods. Briefly, samples were run on 10% SDS-PAGE gels, and transferred onto nitrocellulose membranes (Amersham Protran 0.2μm NC; GE Healthcare Life Sciences). In some cases, blots were cut horizontally prior to primary antibody incubation to allow for the assessment of two different proteins. Blots were washed three times in 1X Tris-Buffered Saline (TBS) and once in 1X TBS with 1% Tween 20 (TBST) for 10 min each. The following antibodies and concentrations were used: rabbit anti-p23 at 1:50,000, rabbit anti-Akr1B-2 at 1:100,000 (produced to full-length *Drosophila* p23 (Q9VH95) by GenScript) and mouse anti-α-Tubulin at 1:5000 (AB_477593; Sigma-Aldrich). Antibodies were diluted in western blot block (5% non-fat dry milk in TBST) and incubated over-night at 4°C. Blots were washed four times with TBS and once with TBST, for 10 min each. The following secondaries were used at 1:5000 in 5mL western blot block: Peroxidase-AffiniPure Goat Anti Rabbit IgG (H+L) and Peroxidase-AffiniPure Goat Anti Mouse IgG (H+L) (Jackson ImmunoResearch Laboratories). Blots were washed four times with TBS and twice with TBST, for 10 min each. Blots were developed with SuperSignal West Pico Chemiluminescent Substrate or SuperSignal West Femto Maximum Sensitivity Substrate (SCR_008452; Thermo Scientific). Blots were imaged and analyzed on the Amersham Imager 600 series Scanning Chemiluminescence (GE Healthcare Life Sciences). Data were collected and analyzed in Excel and graphs were generated and statistical analyses performed in Prism.

## Supplemental Information

**Supplemental Figure 1:**
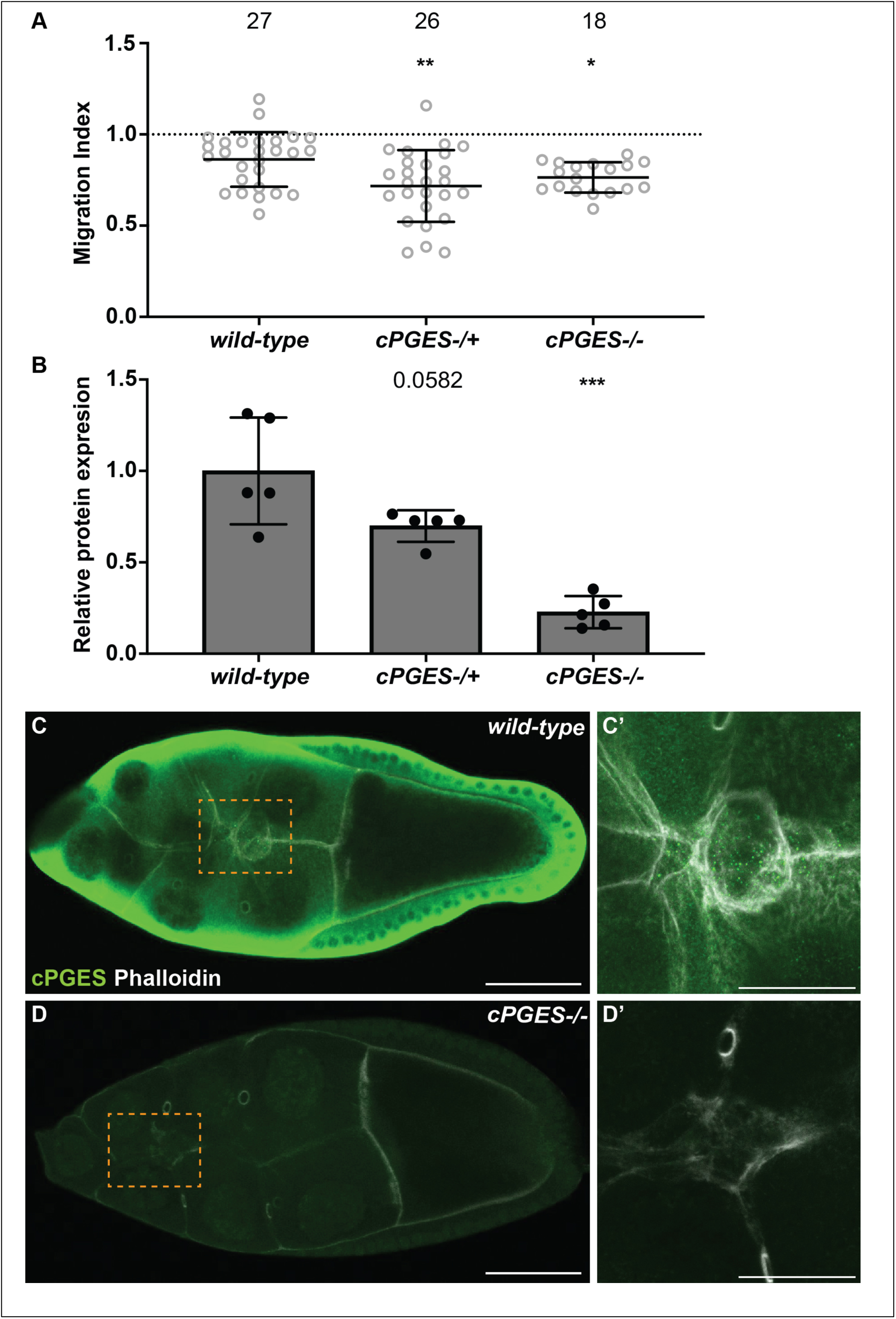
Heterozygosity for *cPGES* delays border cell migration and cPGES is ubiquitously expressed at S9. A. Graph of migration index for the indicated genotypes. Each circle represents a single follicle; n = number of follicles. The dotted line indicates on-time border cell migration, lines = averages, error bars = SD., and * p<0.05 and ** p<0.01, unpaired t-test, two-tailed. B. Graph quantifying cPGES protein levels by western blot analyses normalized to wild-type levels; loading was normalized based on α-Tubulin levels. C-D’. Maximum projections of 3 confocal slices of S9 follicles stained for cPGES (green) and F-actin (phalloidin, white) for the indicated genotypes; higher resolution images of the border cells (orange dashed boxes in C and D) shown in C’ and D’. Images brightened by 30% to increase clarity. Scale bars = 50μm for C, D and 25μm C’, D’. Genotypes used: *wild-type* (*yw*), *cPGES-/+* (*cPGES^EY05607^/+*), and *cPGES-/-* (*cPGES^EY05607/^*^EY05607^). Heterozygosity for the mutation in *cPGES* delays border cell migration to a similar extent as the homozygotes (A). *cPGES-/+* causes a 30% reduction in protein level, while *cPGES-/-* results in a 77% reduction. cPGES is expressed in all the cells of wild-type S9 follicle, including the nurse cell substrate and the border cells (C-C’), and cPGES staining is strongly reduced in the *cPGES* mutant (D-D’).

**Supplemental Figure 2:**
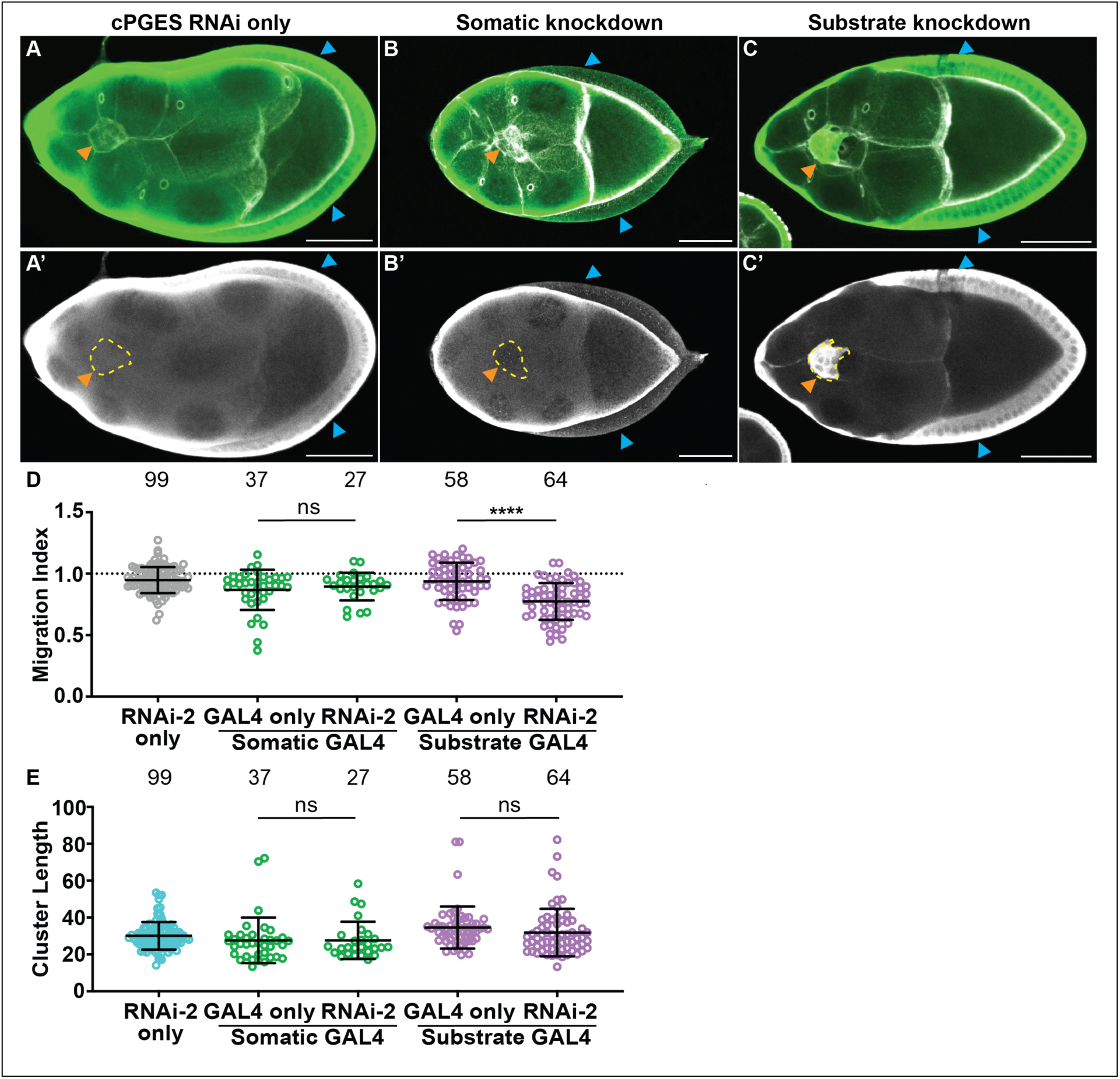
RNAi knockdown of cPGES. A-C. Maximum projections of 3 confocal slices of S9 follicles stained for cPGES (green in merge) and F-actin (phalloidin, white in merge) for the following genotypes: A-A’. cPGES RNAi control (*cPGES RNAi/+*). B-B’. Somatic knockdown of cPGES (*c355 GAL4/+; cPGES RNAi/+*). C-C’. Substrate knockdown of cPGES (*osk GAL4/cPGES RNAi*). cPGES RNAi used in A-C was HMJ24151. Orange arrowheads indicate the border cell cluster, blue arrowheads indicate the outer follicle cells, and yellow dashed lines indicate the position of the outer follicle cells. Images brightened by 30% to increase clarity. Scale bars = 50μm D-E. Graphs of migration index (D) and border cell cluster length (E) for the indicated genotypes. Genotypes include: cPGES RNAi-2 control (*cPGES RNAi-2/+*); somatic GAL4 only (*c355 GAL4/+*); somatic knockdown of cPGES (*c355 GAL4/+; cPGES RNAi-2/+* or *c355 GAL4/+; UAS dicer/+;cPGES RNAi-2/+*), the results were not different with or without UAS Dicer; substrate GAL4 only (*osk GAL4/+*) substrate knockdown of cPGES (*osk GAL4/+;cPGES RNAi-2/+*). cPGES RNAi-2 was GL01292. Each circle represents a single follicle; n = number of follicles. In D, the dotted line indicates on-time border cell migration. For D-E, lines = averages and error bars = SD. ns>0.05, **** p<0.0001, unpaired t-test, two-tailed. cPGES is expressed in all cells of the S9 follicle, including the border cells (A). Whereas, cPGES is expressed only in the substrate in the somatic knockdown (B), and in the somatic cells in the substrate knockdown (C). The second cPGES RNAi line recapitulates what was observed in Fig. 2, as somatic knockdown of cPGES with RNAi-2 exhibits on-time border cell migration, whereas substrate knockdown delays migration (D). Border cell cluster length is unaffected by knockdown of cPGES with RNAi-2 in either the somatic cells or the substrate (E).

**Supplemental Figure 3:**
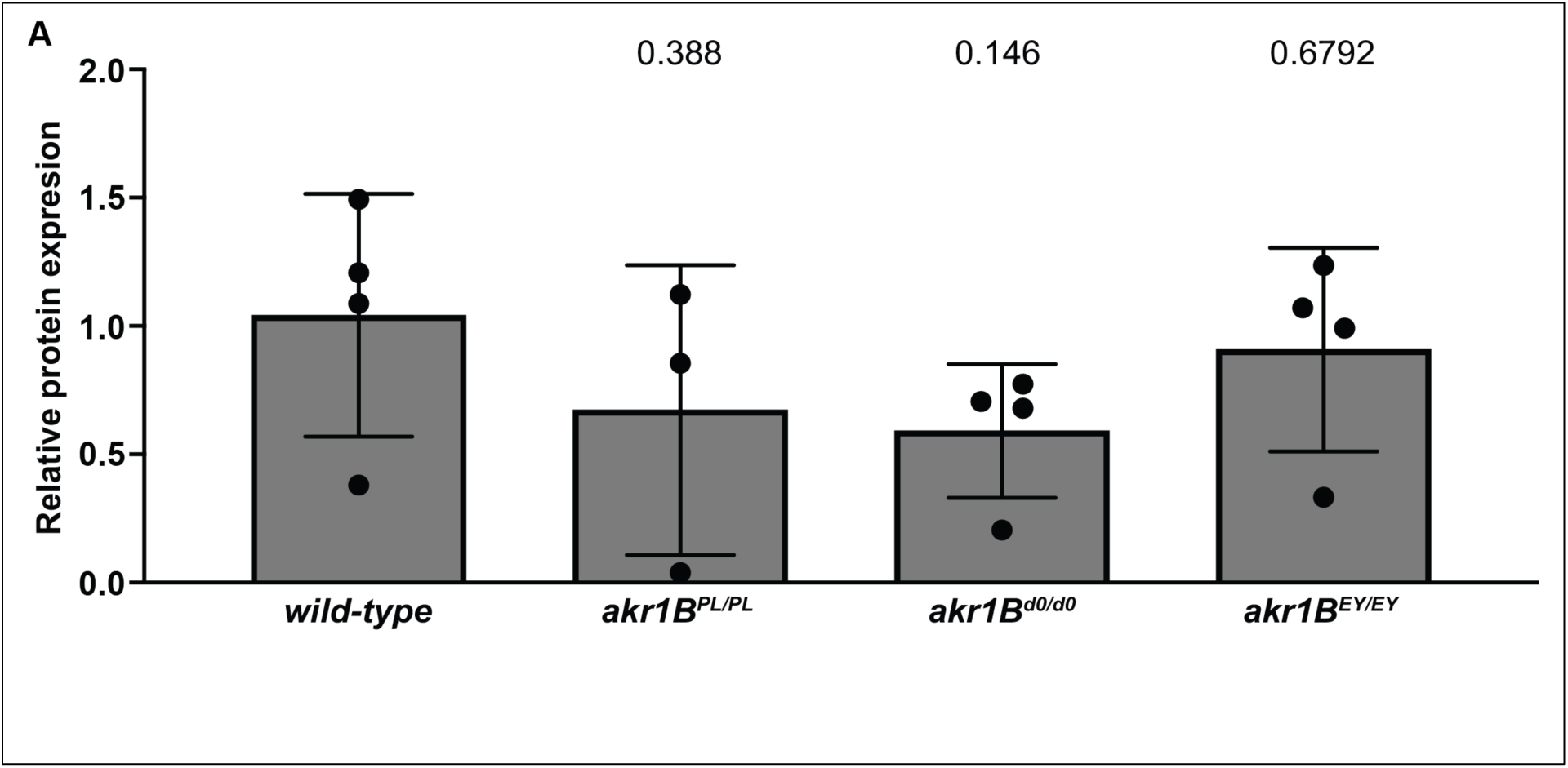
*akr1B* alleles mildly reduce protein level. Graph quantifying protein levels by western blot analyses for Akr1B relative to wild-type values; the α-Tubulin loading control was used to normalize protein levels. Genotypes: *wild-type* (*yw*)*, akr1B^PL00034/PL00034^, akr1B^d00405/d00405^*, and *akr1B^EY07011/EY07011^*. The *akr1B^EY07011/EY07011^*mutants only exhibit a 9% reduction in protein, whereas *akr1B^PL00034/PL00034^*and *akr1B^d00405/d00405^* reduce Akr1B levels by 33% and 40% compared to *wild-type*.

**Supplemental Figure 4:**
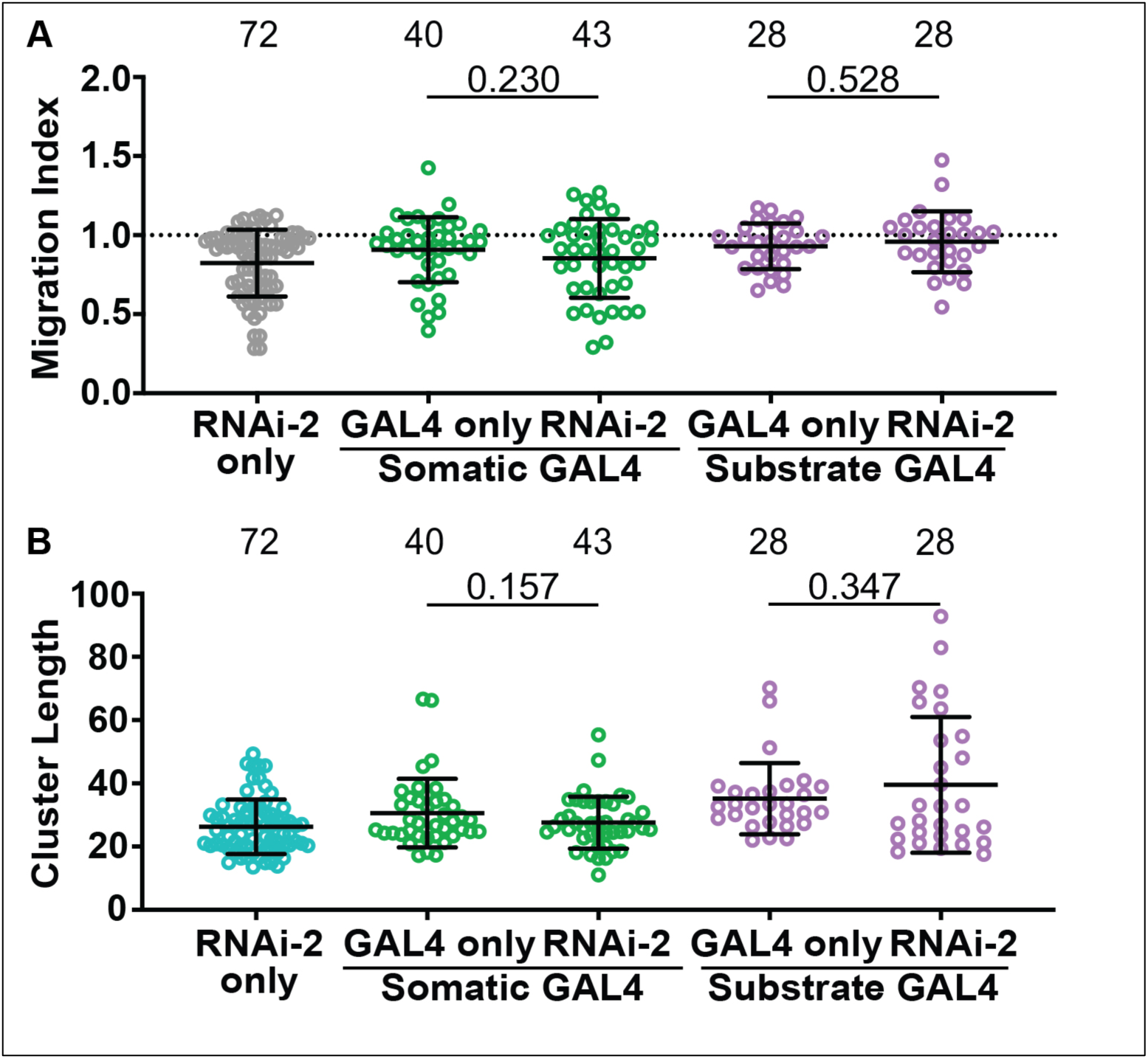
Second Akr1B RNAi line does not affect border cell migration or cluster morphology. A-B. Graphs of migration index (A) and border cell cluster length (B) for the following genotypes include: Akr1B RNAi-2 only control (*akr1B RNAi-2/+*), somatic GAL4 only control (*c355 GAL4/+*), somatic knockdown of Akr1B (*c355 GAL4/+; akr1B RNAi-2/+*), substrate GAL4 only control (*osk GAL4/+*), and substrate knockdown of Akr1B (*osk GAL4/akr1B RNAi-2*). Akr1B RNAi-2 is HMC05226. Each circle represents a single follicle; n = number of follicles. In A, the dotted line indicates on-time border cell migration. For A-B, lines = averages and error bars = SD. p values shown, unpaired t-test, two-tailed. Like the controls, both substrate and somatic knockdown of Akr1B with RNAi-2 exhibits on-time migration (A) and normal cluster length (B).

**Supplemental Figure 5:**
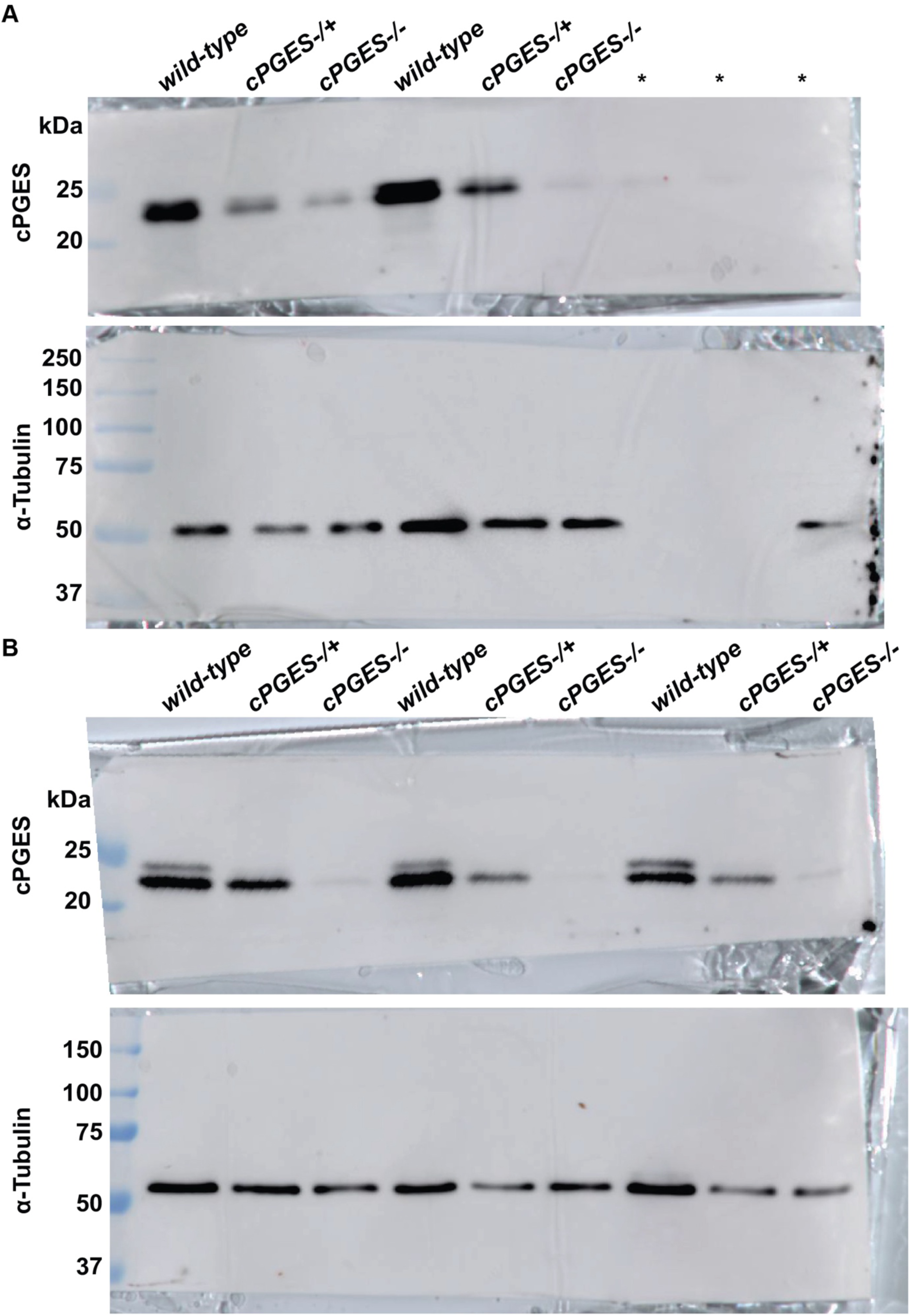
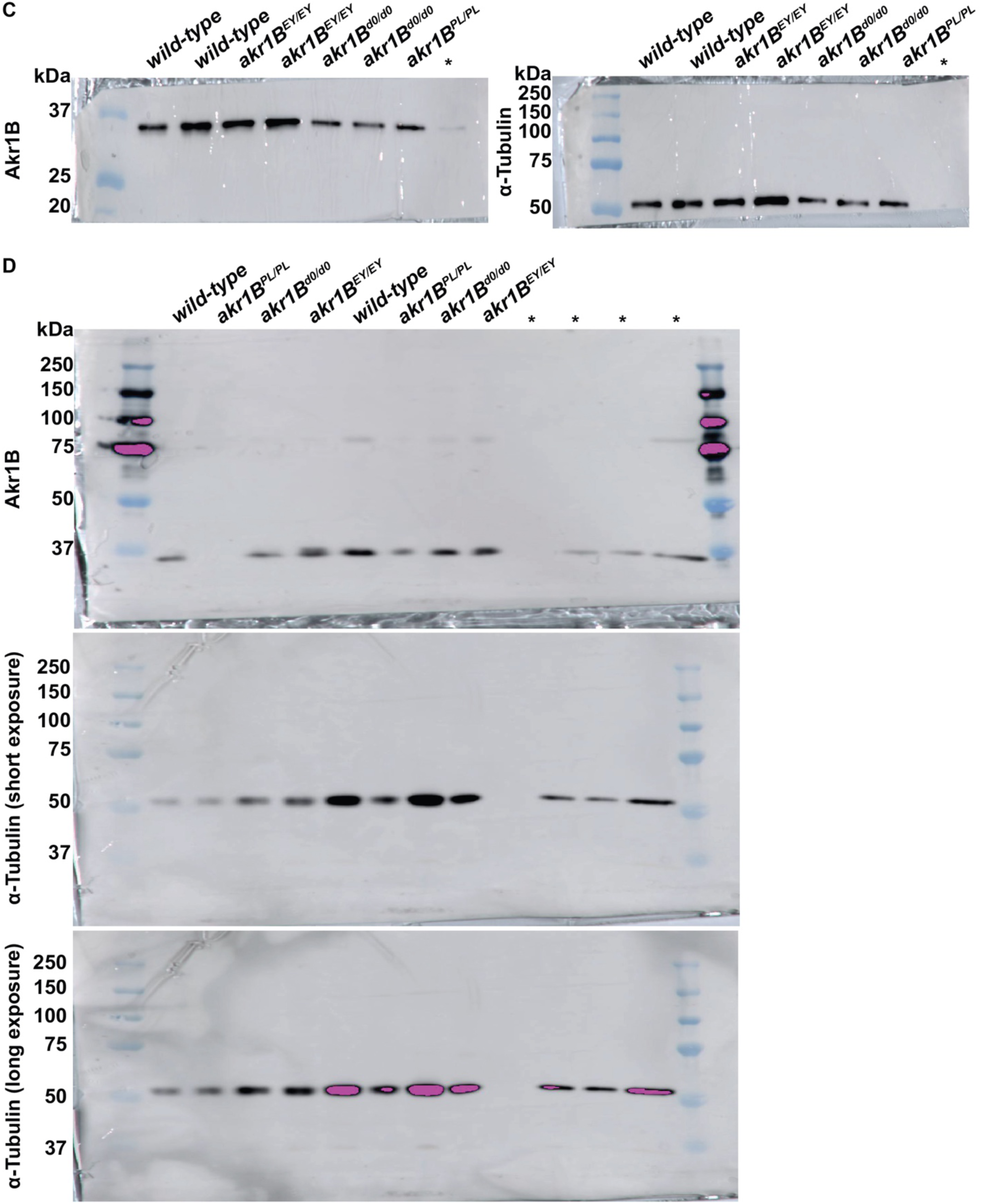
Western blot images. A-D. Images of full western blots used for analyses in SFig. 1 and SFg. 3. Note, lanes that are not used for analysis are labeled with a *. A-B. Western blot for cPGES and α-Tubulin (loading control) for the following genotypes: *wild-type* (*yw*)*, cPGES-/+* (*cPGES^EY05607^/+*)*, and cPGES-/-* (*cPGES^EY05607/EY05607^*)*. cPGES-/+* was either over a *wild-type* or a balancer chromosome. Blot was cut horizontally at ∼35kDa to blot for both proteins at the same time. C-D. Western blots for Akr1B and α-Tubulin (loading control) for the following genotypes: *wild-type* (*yw*)*, akr1B^PL00034/PL00034^, akr1B^d00405/d00405^, akr1B^EY07011/EY07011^*. In C, the blot was cut horizontally at ∼45kDa to blot for both proteins at the same time. In D, the blot was used to detect Akr1B first, then stripped and reprobed to detect α-Tubulin; two different exposures are shown to have the correct exposures for the individual experimental set.

**Table S1. Key Resources Table**

Table of all the reagents and resources used throughout the manuscript.

**Table S2. Genotype by figures**

List of genotypes used in each figure.

Raw data used for quantifications in all the primary and supplemental figures.

**Table S3. Raw data**

Raw data used for quantifications in all the primary and supplemental figures.

## References

Abramoff, MD, Magalhaes, PJ and Ram, SJ (2004) ‘Image processing with ImageJ’, Biophotonics Int. 11: 36–42.

Aguilar-Cuenca, R., Juanes-Garcia, A. and Vicente-Manzanares, M. (2014) ‘Myosin II in mechanotransduction: master and commander of cell migration, morphogenesis, and cancer’, Cell Mol Life Sci 71(3): 479–92.

Alcolea, S., Anton, R., Camacho, M., Soler, M., Alfranca, A., Aviles-Jurado, F. X., Redondo, J. M., Quer, M., Leon, X. and Vila, L. (2012) ‘Interaction between head and neck squamous cell carcinoma cells and fibroblasts in the biosynthesis of PGE2’, J Lipid Res 53(4): 630–42.

Ansari, H. R., Kaddour-Djebbar, I. and Abdel-Latif, A. A. (2004) ‘Effects of prostaglandin F2alpha, latanoprost and carbachol on phosphoinositide turnover, MAP kinases, myosin light chain phosphorylation and contraction and functional existence and expression of FP receptors in bovine iris sphincter’, Exp Eye Res 78(2): 285–96.

Aranjuez, G., Burtscher, A., Sawant, K., Majumder, P. and McDonald, J. A. (2016) ‘Dynamic myosin activation promotes collective morphology and migration by locally balancing oppositional forces from surrounding tissue’, Mol Biol Cell 27(12): 1898–910.

Arnould, T., Thibaut-Vercruyssen, R., Bouaziz, N., Dieu, M., Remacle, J. and Michiels, C. (2001) ‘PGF(2alpha), a prostanoid released by endothelial cells activated by hypoxia, is a chemoattractant candidate for neutrophil recruitment’, Am J Pathol 159(1): 345–57.

Bai, X. M., Zhang, W., Liu, N. B., Jiang, H., Lou, K. X., Peng, T., Ma, J., Zhang, L., Zhang, H. and Leng, J. (2009) ‘Focal adhesion kinase: important to prostaglandin E2-mediated adhesion, migration and invasion in hepatocellular carcinoma cells’, Oncol Rep 21(1): 129–36.

Banerjee, S. (2021) ‘Aldo Keto Reductases AKR1B1 and AKR1B10 in Cancer: Molecular Mechanisms and Signaling Networks’, Adv Exp Med Biol 1347: 65–82.

Barriga, E. H., Franze, K., Charras, G. and Mayor, R. (2018) ‘Tissue stiffening coordinates morphogenesis by triggering collective cell migration in vivo’, Nature 554(7693): 523–527.

Butcher, D. T., Alliston, T. and Weaver, V. M. (2009) ‘A tense situation: forcing tumour progression’, Nat Rev Cancer 9(2): 108–22.

Cai, D., Chen, S. C., Prasad, M., He, L., Wang, X., Choesmel-Cadamuro, V., Sawyer, J. K., Danuser, G. and Montell, D. J. (2014) ‘Mechanical feedback through E-cadherin promotes direction sensing during collective cell migration’, Cell 157(5): 1146–59.

Cano, L. Q., Lavery, D. N., Sin, S., Spanjaard, E., Brooke, G. N., Tilman, J. D., Abroaf, A., Gaughan, L., Robson, C. N., Heer, R. et al. (2015) ‘The co-chaperone p23 promotes prostate cancer motility and metastasis’, Mol Oncol 9(1): 295–308.

Cha, Y. I., Kim, S. H., Sepich, D., Buchanan, F. G., Solnica-Krezel, L. and DuBois, R. N. (2006) ‘Cyclooxygenase-1-derived PGE2 promotes cell motility via the G-protein-coupled EP4 receptor during vertebrate gastrulation’, Genes Dev 20(1): 77–86.

Cha, Y. I., Kim, S. H., Solnica-Krezel, L. and Dubois, R. N. (2005) ‘Cyclooxygenase-1 signaling is required for vascular tube formation during development’, Dev Biol 282(1): 274–83.

Chandler, C., Liu, T., Buckanovich, R. and Coffman, L. G. (2019) ‘The double edge sword of fibrosis in cancer’, Transl Res 209: 55–67.

Cox, T. R. and Erler, J. T. (2014) ‘Molecular pathways: connecting fibrosis and solid tumor metastasis’, Clin Cancer Res 20(14): 3637–43.

Danny L. Brower, Michael Wilcox, Michel Piovant, Richard J. Smith, and Lorrel A. Reger (1984) ‘Relatedcell-surfaceantigensexpressedwithpositionalspecificityin Drosophila imaginal discs’, Developmental Biology 81: 7485–7489.

DeLuca, S. Z. and Spradling, A. C. (2018) ‘Efficient Expression of Genes in the Drosophila Germline Using a UAS Promoter Free of Interference by Hsp70 piRNAs’, Genetics 209(2): 381–387.

Demirkol Canli, S., Seza, E. G., Sheraj, I., Gomceli, I., Turhan, N., Carberry, S., Prehn, J. H. M., Gure, A. O. and Banerjee, S. (2020) ‘Evaluation of an aldo-keto reductase gene signature with prognostic significance in colon cancer via activation of epithelial to mesenchymal transition and the p70S6K pathway’, Carcinogenesis 41(9): 1219–1228.

Diao, Ge, Huang, Jie, Zheng, Xiuhui, Sun, Xinwei, Tian, Min, Han, Jian and Guo, Jianxin (2021) ‘Prostaglandin E2 serves a dual role in regulating the migration of dendritic cells’, Int J Mol Med 47(1): 207–218.

Dinkins, M. B., Fratto, V. M. and Lemosy, E. K. (2008) ‘Integrin alpha chains exhibit distinct temporal and spatial localization patterns in epithelial cells of the Drosophila ovary’, Dev Dyn 237(12): 3927–39.

Eble, J. A. and Niland, S. (2019) ‘The extracellular matrix in tumor progression and metastasis’, Clin Exp Metastasis 36(3): 171–198.

Elwakeel, E., Brune, B. and Weigert, A. (2019) ‘PGE(2) in fibrosis and cancer: Insights into fibroblast activation’, Prostaglandins Other Lipid Mediat 143: 106339.

Enyedi, B., Jelcic, M. and Niethammer, P. (2016) ‘The Cell Nucleus Serves as a Mechanotransducer of Tissue Damage-Induced Inflammation’, Cell 165(5): 1160–1170.

Fife, C. M., McCarroll, J. A. and Kavallaris, M. (2014) ‘Movers and shakers: cell cytoskeleton in cancer metastasis’, Br J Pharmacol 171(24): 5507–23.

Fox, E. F., Lamb, M. C., Mellentine, S. Q. and Tootle, T. L. (2020) ‘Prostaglandins regulate invasive, collective border cell migration’, Mol Biol Cell 31(15): 1584–1594.

Friedl, P. and Gilmour, D. (2009) ‘Collective cell migration in morphogenesis, regeneration and cancer’, Nat Rev Mol Cell Biol 10(7): 445–57.

Fujino, H., Toyomura, K., Chen, X. B., Regan, J. W. and Murayama, T. (2011) ‘Prostaglandin E(2) regulates cellular migration via induction of vascular endothelial growth factor receptor-1 in HCA-7 human colon cancer cells’, Biochem Pharmacol 81(3): 379–87.

Funk, C. D. (2001) ‘Prostaglandins and leukotrienes: advances in eicosanoid biology’, Science 294(5548): 1871–5.

Gasparski, A. N., Ozarkar, S. and Beningo, K. A. (2017) ‘Transient mechanical strain promotes the maturation of invadopodia and enhances cancer cell invasion in vitro’, J Cell Sci 130(11): 1965–1978.

Giedt, M. S. and Tootle, T. L. (2023) ‘The Vast Utility of Drosophila Oogenesis’, Methods Mol Biol 2626: 1–36.

Groen, C. M., Spracklen, A. J., Fagan, T. N. and Tootle, T. L. (2012) ‘Drosophila Fascin is a novel downstream target of prostaglandin signaling during actin remodeling’, Mol Biol Cell 23(23): 4567–78.

Hall, M. S., Alisafaei, F., Ban, E., Feng, X., Hui, C. Y., Shenoy, V. B. and Wu, M. (2016) ‘Fibrous nonlinear elasticity enables positive mechanical feedback between cells and ECMs’, Proc Natl Acad Sci U S A 113(49): 14043–14048.

Jakobsson, P. J., Thoren, S., Morgenstern, R. and Samuelsson, B. (1999) ‘Identification of human prostaglandin E synthase: a microsomal, glutathione-dependent, inducible enzyme, constituting a potential novel drug target’, Proc Natl Acad Sci U S A 96(13): 7220–5.

Jongthawin, J., Chusorn, P., Techasen, A., Loilome, W., Boonmars, T., Thanan, R., Puapairoj, A., Khuntikeo, N., Tassaneeyakul, W., Yongvanit, P. et al. (2014) ‘PGE2 signaling and its biosynthesis-related enzymes in cholangiocarcinoma progression’, Tumour Biol 35(8): 8051–64.

Kai, F., Laklai, H. and Weaver, V. M. (2016) ‘Force Matters: Biomechanical Regulation of Cell Invasion and Migration in Disease’, Trends Cell Biol 26(7): 486–497.

Kamei, D., Murakami, M., Sasaki, Y., Nakatani, Y., Majima, M., Ishikawa, Y., Ishii, T., Uematsu, S., Akira, S., Hara, S. et al. (2009) ‘Microsomal prostaglandin E synthase-1 in both cancer cells and hosts contributes to tumour growth, invasion and metastasis’, Biochem J 425(2): 361–71.

Karavitis, J., Hix, L. M., Shi, Y. H., Schultz, R. F., Khazaie, K. and Zhang, M. (2012) ‘Regulation of COX2 expression in mouse mammary tumor cells controls bone metastasis and PGE2-induction of regulatory T cell migration’, PLoS One 7(9): e46342.

Kelly Cant, Brenda A Knowles, Mark S. Mooseker, and Lynn Cooley (1994) ‘Drosophila Singed, a Fascin Homolog, is Required for Acin Bundel Formation during Oogenesis and Bristle Extension’, Celll Biology 125: 369–380.

Kim, J. I., Lakshmikanthan, V., Frilot, N. and Daaka, Y. (2010) ‘Prostaglandin E2 promotes lung cancer cell migration via EP4-betaArrestin1-c-Src signalsome’, Mol Cancer Res 8(4): 569–77.

Kobayashi, K., Omori, K. and Murata, T. (2018) ‘Role of prostaglandins in tumor microenvironment’, Cancer Metastasis Rev 37(2-3): 347–354.

Lamb, M. C., Anliker, K. K. and Tootle, T. L. (2020) ‘Fascin regulates protrusions and delamination to mediate invasive, collective cell migration in vivo’, Dev Dyn 249(8): 961–982.

Lamb, M. C., Kaluarachchi, C. P., Lansakara, T. I., Mellentine, S. Q., Lan, Y., Tivanski, A. V. and Tootle, T. L. (2021) ‘Fascin limits Myosin activity within Drosophila border cells to control substrate stiffness and promote migration’, Elife 10.

Legler, D. F., Krause, P., Scandella, E., Singer, E. and Groettrup, M. (2006) ‘Prostaglandin E2 is generally required for human dendritic cell migration and exerts its effect via EP2 and EP4 receptors’, J Immunol 176(2): 966–73.

Li, H. J., Reinhardt, F., Herschman, H. R. and Weinberg, R. A. (2012) ‘Cancer-stimulated mesenchymal stem cells create a carcinoma stem cell niche via prostaglandin E2 signaling’, Cancer Discov 2(9): 840–55.

Liu, F., Mih, J. D., Shea, B. S., Kho, A. T., Sharif, A. S., Tager, A. M. and Tschumperlin, D. J. (2010) ‘Feedback amplification of fibrosis through matrix stiffening and COX-2 suppression’, J Cell Biol 190(4): 693–706.

Llense, F. and Martin-Blanco, E. (2008) ‘JNK signaling controls border cell cluster integrity and collective cell migration’, Curr Biol 18(7): 538–44.

Lo, C. M., Wang, H. B., Dembo, M. and Wang, Y. L. (2000) ‘Cell movement is guided by the rigidity of the substrate’, Biophys J 79(1): 144–52.

Lomakin, A. J., Cattin, C. J., Cuvelier, D., Alraies, Z., Molina, M., Nader, G. P. F., Srivastava, N., Saez, P. J., Garcia-Arcos, J. M., Zhitnyak, I. Y. et al. (2020) ‘The nucleus acts as a ruler tailoring cell responses to spatial constraints’, Science 370(6514).

Lu, D., Han, C. and Wu, T. (2012) ‘Microsomal prostaglandin E synthase-1 promotes hepatocarcinogenesis through activation of a novel EGR1/beta-catenin signaling axis’, Oncogene 31(7): 842–57.

Majumder, P., Aranjuez, G., Amick, J. and McDonald, J. A. (2012) ‘Par-1 controls myosin-II activity through myosin phosphatase to regulate border cell migration’, Curr Biol 22(5): 363–72.

Mayoral, R., Fernandez-Martinez, A., Bosca, L. and Martin-Sanz, P. (2005) ‘Prostaglandin E2 promotes migration and adhesion in hepatocellular carcinoma cells’, Carcinogenesis 26(4): 753–61.

Menter, D. G. and Dubois, R. N. (2012) ‘Prostaglandins in cancer cell adhesion, migration, and invasion’, Int J Cell Biol 2012: 723419.

Mohan, K., Luo, T., Robinson, D. N. and Iglesias, P. A. (2015) ‘Cell shape regulation through mechanosensory feedback control’, J R Soc Interface 12(109): 20150512.

Montell, D. J. (2003) ‘Border-cell migration: the race is on’, Nat Rev Mol Cell Biol 4(1): 13–24.

Montell, D. J., Yoon, W. H. and Starz-Gaiano, M. (2012) ‘Group choreography: mechanisms orchestrating the collective movement of border cells’, Nat Rev Mol Cell Biol 13(10): 631–45.

Nakanishi, M., Menoret, A., Tanaka, T., Miyamoto, S., Montrose, D. C., Vella, A. T. and Rosenberg, D. W. (2011) ‘Selective PGE(2) suppression inhibits colon carcinogenesis and modifies local mucosal immunity’, Cancer Prev Res (Phila) 4(8): 1198–208.

Nakanishi, M., Montrose, D. C., Clark, P., Nambiar, P. R., Belinsky, G. S., Claffey, K. P., Xu, D. and Rosenberg, D. W. (2008) ‘Genetic deletion of mPGES-1 suppresses intestinal tumorigenesis’, Cancer Res 68(9): 3251–9.

Oakes, P. W. (2018) ‘Balancing forces in migration’, Curr Opin Cell Biol 54: 43–49.

Osma-Garcia, I. C., Punzon, C., Fresno, M. and Diaz-Munoz, M. D. (2016) ‘Dose-dependent effects of prostaglandin E2 in macrophage adhesion and migration’, Eur J Immunol 46(3): 677–88.

Patel, N. H., Snow, P. M. and Goodman, C. S. (1987) ‘Characterization and cloning of fasciclin III: a glycoprotein expressed on a subset of neurons and axon pathways in Drosophila’, Cell 48(6): 975–88.

Piersma, B., Hayward, M. K. and Weaver, V. M. (2020) ‘Fibrosis and cancer: A strained relationship’, Biochim Biophys Acta Rev Cancer 1873(2): 188356.

Platt, J. L. and Michael, A. F. (1983) ‘Retardation of fading and enhancement of intensity of immunofluorescence by p-phenylenediamine’, J Histochem Cytochem 31(6): 840–2.

Ren, Y., Zhang, Y., Liu, J., Liu, P., Yang, J., Guo, D., Tang, A. and Tao, J. (2021) ‘Matrix hardness regulates the cancer cell malignant progression through cytoskeletal network’, Biochem Biophys Res Commun 541: 95–101.

Rieder, F., Georgieva, M., Schirbel, A., Artinger, M., Zugner, A., Blank, M., Brenmoehl, J., Scholmerich, J. and Rogler, G. (2010) ‘Prostaglandin E2 inhibits migration of colonic lamina propria fibroblasts’, Inflamm Bowel Dis 16(9): 1505–13.

Sales, K. J., Grant, V. and Jabbour, H. N. (2008) ‘Prostaglandin E2 and F2alpha activate the FP receptor and up-regulate cyclooxygenase-2 expression via the cyclic AMP response element’, Mol Cell Endocrinol 285(1-2): 51–61.

Scarpa, E. and Mayor, R. (2016) ‘Collective cell migration in development’, J Cell Biol 212(2): 143–55.

Singh, B., Berry, J. A., Shoher, A., Ramakrishnan, V. and Lucci, A. (2005) ‘COX-2 overexpression increases motility and invasion of breast cancer cells’, Int J Oncol 26(5): 1393–9.

Spracklen, A. J., Kelpsch, D. J., Chen, X., Spracklen, C. N. and Tootle, T. L. (2014) ‘Prostaglandins temporally regulate cytoplasmic actin bundle formation during Drosophila oogenesis’, Mol Biol Cell 25(3): 397–411.

Stamatakis, K., Jimenez-Martinez, M., Jimenez-Segovia, A., Chico-Calero, I., Conde, E., Galan-Martinez, J., Ruiz, J., Pascual, A., Barrocal, B., Lopez-Perez, R. et al. (2015) ‘Prostaglandins induce early growth response 1 transcription factor mediated microsomal prostaglandin E2 synthase up-regulation for colorectal cancer progression’, Oncotarget 6(37): 39941–59.

Stuelten, C. H., Parent, C. A. and Montell, D. J. (2018) ‘Cell motility in cancer invasion and metastasis: insights from simple model organisms’, Nat Rev Cancer 18(5): 296–312.

Tanikawa, N., Ohmiya, Y., Ohkubo, H., Hashimoto, K., Kangawa, K., Kojima, M., Ito, S. and Watanabe, K. (2002) ‘Identification and characterization of a novel type of membrane-associated prostaglandin E synthase’, Biochem Biophys Res Commun 291(4): 884–9.

Tanioka, T., Nakatani, Y., Semmyo, N., Murakami, M. and Kudo, I. (2000) ‘Molecular identification of cytosolic prostaglandin E2 synthase that is functionally coupled with cyclooxygenase-1 in immediate prostaglandin E2 biosynthesis’, J Biol Chem 275(42): 32775–82.

Telley, I. A., Gaspar, I., Ephrussi, A. and Surrey, T. (2012) ‘Aster migration determines the length scale of nuclear separation in the Drosophila syncytial embryo’, J Cell Biol 197(7): 887–95.

Thibault, S. T., Singer, M. A., Miyazaki, W. Y., Milash, B., Dompe, N. A., Singh, C. M., Buchholz, R., Demsky, M., Fawcett, R., Francis-Lang, H. L. et al. (2004) ‘A complementary transposon tool kit for Drosophila melanogaster using P and piggyBac’, Nat Genet 36(3): 283–7.

Tootle, T. L. (2013) ‘Genetic insights into the in vivo functions of prostaglandin signaling’, Int J Biochem Cell Biol 45(8): 1629–32.

Tootle, T. L. and Spradling, A. C. (2008) ‘Drosophila Pxt: a cyclooxygenase-like facilitator of follicle maturation’, Development 135(5): 839–47.

van Helden, S. F., Oud, M. M., Joosten, B., Peterse, N., Figdor, C. G. and van Leeuwen, F. N. (2008) ‘PGE2-mediated podosome loss in dendritic cells is dependent on actomyosin contraction downstream of the RhoA-Rho-kinase axis’, J Cell Sci 121(Pt 7): 1096–106.

van Helvert, S. and Friedl, P. (2016) ‘Strain Stiffening of Fibrillar Collagen during Individual and Collective Cell Migration Identified by AFM Nanoindentation’, ACS Appl Mater Interfaces 8(34): 21946–55.

van Helvert, S., Storm, C. and Friedl, P. (2018) ‘Mechanoreciprocity in cell migration’, Nat Cell Biol 20(1): 8–20.

Vicente-Manzanares, M., Ma, X., Adelstein, R. S. and Horwitz, A. R. (2009) ‘Non-muscle myosin II takes centre stage in cell adhesion and migration’, Nat Rev Mol Cell Biol 10(11): 778–90.

Vo, B. T., Morton, D., Jr., Komaragiri, S., Millena, A. C., Leath, C. and Khan, S. A. (2013) ‘TGF-beta effects on prostate cancer cell migration and invasion are mediated by PGE2 through activation of PI3K/AKT/mTOR pathway’, Endocrinology 154(5): 1768–79.

Wallace, A. E., Sales, K. J., Catalano, R. D., Anderson, R. A., Williams, A. R., Wilson, M. R., Schwarze, J., Wang, H., Rossi, A. G. and Jabbour, H. N. (2009) ‘Prostaglandin F2alpha-F-prostanoid receptor signaling promotes neutrophil chemotaxis via chemokine (C-X-C motif) ligand 1 in endometrial adenocarcinoma’, Cancer Res 69(14): 5726–33.

Wolf, K., Wu, Y. I., Liu, Y., Geiger, J., Tam, E., Overall, C., Stack, M. S. and Friedl, P. (2007) ‘Multi-step pericellular proteolysis controls the transition from individual to collective cancer cell invasion’, Nat Cell Biol 9(8): 893–904.

Wu, X., Li, X., Fu, Q., Cao, Q., Chen, X., Wang, M., Yu, J., Long, J., Yao, J., Liu, H. et al. (2017) ‘AKR1B1 promotes basal-like breast cancer progression by a positive feedback loop that activates the EMT program’, J Exp Med 214(4): 1065–1079.

Xu, C., You, X., Liu, W., Sun, Q., Ding, X., Huang, Y. and Ni, X. (2015) ‘Prostaglandin F2alpha regulates the expression of uterine activation proteins via multiple signalling pathways’, Reproduction 149(1): 139–46.

Zaccai, M. and Lipshitz, H. D. (1996) ‘Differential distributions of two adducin-like protein isoforms in the Drosophila ovary and early embryo’, Zygote 4(2): 159–66.

